# Closing The Loop: A Dynamic Neural Network Model Integrating Decision making and Metacognition

**DOI:** 10.1101/2025.03.06.641797

**Authors:** Weiwen Lu, Xiaohong Wan

## Abstract

Decision making under uncertainty entails selecting optimal actions from noisy evidence. Currently influential models, such as drift-diffusion and attractor frameworks, posit decisions as bottom-up stochastic evidence accumulation from sensory inputs but ignore critical interactions between decision, commitment, and metacognition. While these models explain basic choice behaviors and accompanying confidence, they fail to reconcile many empirical findings, including motor-area encoding of decision variables and time-dependent urgency signals. We present a closed-loop neural network model unifying three modules: a decision module accumulating evidence, a motor module implementing action thresholds, and a metacognition module regulating deliberation through dual feedback pathways to suppress noise-driven errors and accelerate decision commitment under time constraints, respectively. This architecture can account for crucial characteristics of decision-making and metacognition that were empirically observed. By integrating decision, motor, and metacognitive dynamics, our model provides a biologically grounded framework for optimal decision making, offering testable predictions for neural and behavioral studies.

## Introduction

Decision making is a dynamic cognitive process that transforms sensory inputs into adaptive actions through sequential integration of evidence, evaluation of alternatives, and commitment to a choice (*1–3*). The currently influential decision-making models, such as the drift-diffusion model (DDM; 2, 4, 5) and the neural network attractor model (NNAM; 3, 6, 7), however, conceptualize the decision-making process primarily as bottom-up stochastic evidence accumulation over time until a specified threshold is reached. Once the threshold is crossed, the decision process is terminated, and an action response is elicited. These simplified models successfully account for various aspects of decision-making behavior, such as choices and response times (RTs), and also provide insights into the neural activities responsible for encoding decision variables associated with evidence accumulation (*2*, *3*).

Decision making under uncertainty is accompanied by metacognition, which involves in assessing the quality of decisions to form beliefs about their correctness or superiority compared to alternative options, referred to as confidence (*8*). The above decision-making models have been also expanded to incorporate confidence-related behavior (*9–11*). These models suggest that confidence is a direct consequence of decision making and thus determined by the strength or quantity of the same decision variable that is accumulated through the decision-making process. This view highlights a common neural mechanism shared between decision making and metacognition.

However, empirical studies present mounting evidence that decision making encompasses much more complexity than what can be addressed by these simple models, particularly in the following three aspects. Firstly, motor regions also play a significant role in the decision-making process, rather than merely execute action responses following decisions. The primary motor cortex (M1) (*12*), the premotor cortex (*12*), and the pre-supplementary motor area (pre-SMA, ref. *13*), exhibit neural activities that encode quantitative decision variables, similar to the brain regions typically associated with decision making, such as the lateral intraparietal area (LIP, ref. *2*). Critically, the quantified momentary neural activities in the motor areas precisely calibrate decision accuracy (*12*).

Secondly, in the classical decision-making models, there are often instances where the predefined threshold cannot be reached within a reasonable time interval, leading to incomplete decisions, particularly under weak evidence. However, the occurrence of missing choices is much less frequent in real animal and human decision making even under uncertainty. In these instances, an additional signal, independent of the process of evidence accumulation, is found to enhance neural activities to expedite the decision-making process towards reaching the threshold (*14*, *15*), likened to a reduction in the threshold required to terminate a decision (collapsing boundary). The presence of this additional signal, known as the urgency signal, is proportional to the elapsed time during decision making (*14*, *15*). The urgency signal has been theoretically and empirically identified as a crucial element for optimal decision making (*5*, *16*). Nevertheless, how the urgency signal emerges thus far remain unresolved (*17*, *18*).

Thirdly, confidence has been found to be behaviorally dissociable from actual accuracy in decision making (*19–21*). Further, the brain regions responsible for coding confidence, such as the dorsal anterior cingulate cortex (dACC), are distinct from those involved in the decision-making processes (*22*, *23*). These findings suggest that the generation of confidence involves additional components that operate independently of the decision-making process (*24*, *25*). Importantly, metacognition is not merely a post-hoc evaluation but actively optimizes decisions through real-time feedback, particularly in uncertain environments (26, 27). However, the currently prevailing decision-making models (*1–11*, but see *24*, *25*), which lack explicit mechanisms for such top-down control, fail to explain how confidence modulates the decision-making process.

To address these gaps, we present a dynamic neural network model that unifies decision making, motor preparation, and metacognition into a closed-loop system (**Fig. 1**). In this framework, the decision module and motor module mutually inhibit one another, with the motor module hosting decision thresholds to reflect its role in action commitment. The metacognition module monitors uncertainty through feedforward signals from the decision module, generating two regulatory signals onto the decision module and motor module, respectively. By integrating decision, motor, and metacognitive processes together, our model offers a biologically plausible account of adaptive decision making under uncertainty. It not only reconciles conflicting empirical findings but also generates testable predictions for how neural circuits balance speed, accuracy, and confidence in empirical studies.

**Figure 1.**
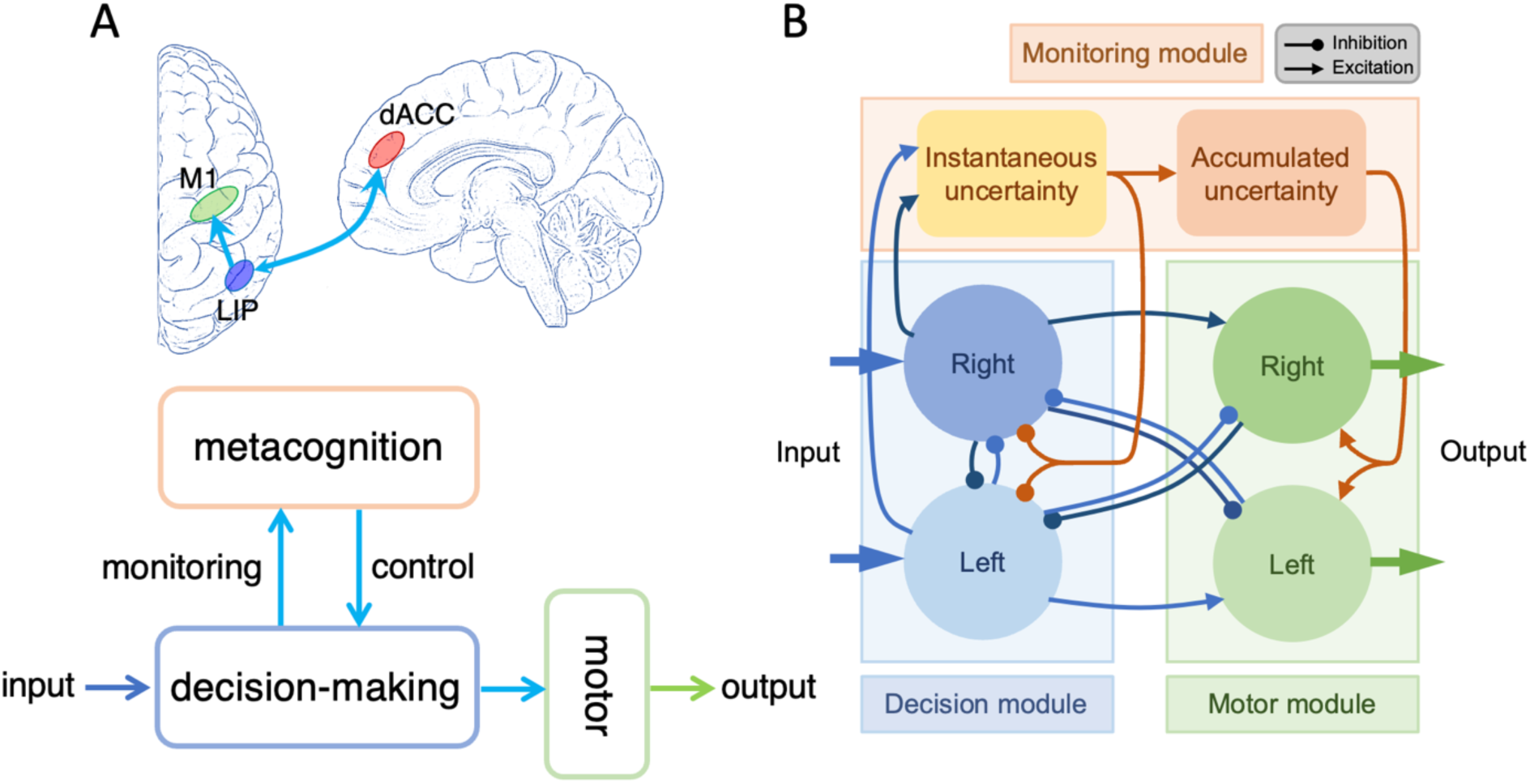
The neural network architecture for decision making and metacognition. (**A**) The currently conceptual and neural models for decision making and metacognition, in which decision-making, motor action and metacognition are separated. (**B**) The closed-loop neural network model comprising of the decision module, the motor module and the metacognition module. Except for the feedforward connections between these modules, as like in the traditional view (**A**), there exist feedbacks from the motor module to the decision module, as well as from the metacognition module to the decision module and the motor module.

## Results

### The neural-network architecture

The proposed neural network model features three interconnected modules (**Fig. 1B**): (1) a decision module that accumulates evidence through competing neuronal populations, as in the classical NNAM model; (2) a motor module that hosts decision thresholds and mutually inhibits the decision module to segregate early and late decision-making phases. It plays a crucial role in translating decisions into actions; and (3) a metacognition module that monitors decision uncertainty via feedforward inputs and regulates the system through dual feedback mechanisms. Instantaneous uncertainty, derived from ongoing decision dynamics and monitoring instantaneous contradictions, suppresses noise-induced activities in the decision module via inhibitory feedback (**Fig. 2D**), while accumulated uncertainty reflecting accumulated contradictions along the decision-making progress generates urgency signals that excitatorily bias the motor module to enforce timely commitments (**Fig. 2E**). By dynamically balancing deliberation and urgency, the model optimizes speed-accuracy tradeoffs and explains how urgency signals emerge, and motor areas affect the decision-making process. Notably, each module represents a unique functional unit that may biologically encompass multiple brain areas, rather than being confined to a single brain area.

**Figure 2.**
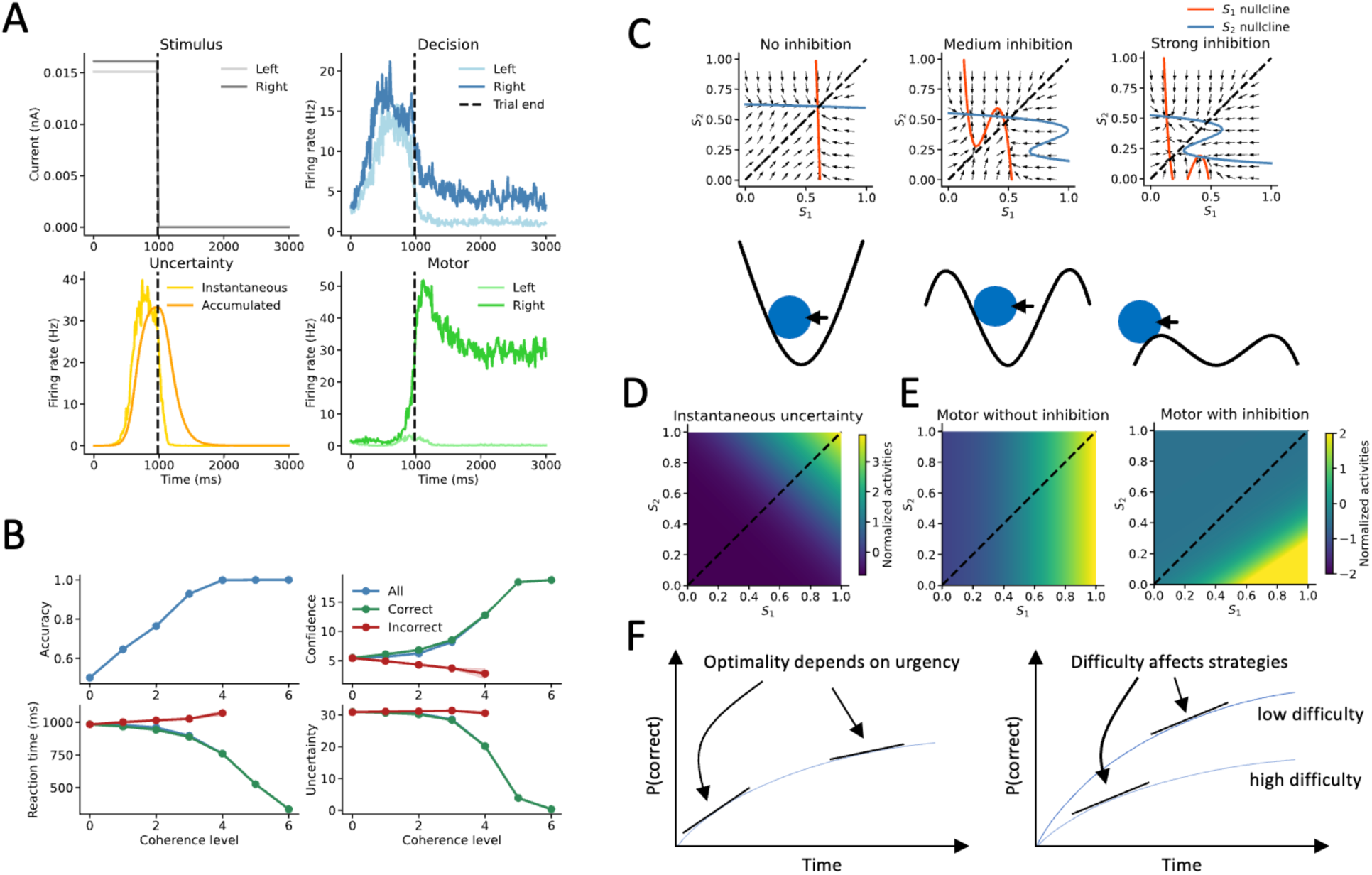
The neural dynamics of the neural network model. (**A**) An instance of neural dynamics of the neural network model. (**B**) The replicates of basic features in decision making and metacognition. (**C**) The illustrations of states under different strengths of mutual inhibitions. (**D**) the instantaneous uncertainty as functions of the states of the decision module. (**E**) The motor activities dependent on the states of the decision module with/wo inter-hemispherical inhibitions between the motor module and the decision module. (**F**) Optimal decision making in trade-offs between accuracy and time.

Computational efficiencies shape cognitive processes. During decision making, a longer time spent on deliberation tends to improve accuracy. However, the incremental gain in decision accuracy diminishes gradually over time (**Fig. 2F**). On the other hand, dedicating excessive time to the ongoing decision-making process incurs a cost of missing out on other potential opportunities. Therefore, achieving optimal decisions requires a strategy balancing the inherent benefits and costs involved in the current decision making (*16*, *28*).

In this model, multiple aspects of the neural network architecture are motivated to achieve optimal decisions. Firstly, in the original NNAM model, a strong mutual inhibition often exists between the two competitive populations of neurons involved in accumulating alternative evidence, easily leading to winner-takes-all (WTA) states, in which one is dominant. This facilitates the reaching of the decision threshold. However, this fast dynamics is vulnerable to noise. By contrast, a weak mutual inhibition in the decision module of the current model reduces the likelihood of early WTA formation (**Fig 2C**). Consequently, a state of high instantaneous uncertainty emerges, which, in turn, suppresses the neural activities of the decision module through inhibitory feedback (**Fig 2D**). This adaptive neural network with negative feedback significantly mitigates the influence of noise, prevents decision states from becoming overly rigid and resistant to changes and allows for further deliberation with insufficient evidence, particularly during the early phase of decision making.

Secondly, the presence of mutual contralateral inhibitory connections between the decision module and the motor module allows for a clear separation of their unique roles in evidence accumulation. The contralateral decision-motor inhibition effectively slows down decision-making as the motor module could only be activated with a certain level of difference between the nodes of the decision module (**Fig. 2E**). Thus, the motor module plays a more significant role during the late phase for stabilizing choice commitments, but has a lesser influence during the early phase of decision making in deliberation.

Lastly, and of utmost importance, the accumulated uncertainty provides excitatory feedback to the neurons in the motor module, serving as the source of the urgency signal. Schematically, a lower rate of evidence accumulation for the dominant option corresponds to a higher level of accumulated uncertainty. This accumulated uncertainty is then transformed into an urgency signal added onto the decision variables accumulated in the motor module, facilitating a timely termination of the decision-making process, even in the presence of uncertainty (**Fig. 2F**). Hence, in the face of uncertainty, the adaptive inhibition from instantaneuous uncertainty and the contralateral decision-motor inhibition serves to support further deliberation over time. On the other hand, the urgency signal from the accumulated uncertainty serves to terminate decision-making timely. These two contradictive effects of uncertainty stem from two different components of uncertainty embedded in the neural circuit.

By synthesizing these dynamic features of the neural network architecture, the currently proposed closed-loop neural network model ensures efficient adaptability of decision states, enabling the achievement of optimal decisions even in the presence of noise and uncertainty.

It is noteworthy that the model employs two complementary variables to evaluate the belief regarding decision correctness during the decision-making process. One variable, serving as a proxy for confidence, is computed by the absolute difference between the neural activities of competing neuron populations in the decision module (‘balance of evidence’, ref. *11*, *29*, *30*).

However, it is important to acknowledge that the current model lacks a specified neural circuit for computing this variable. By contrast, another variable captures the observable state of accumulated uncertainty, which may correspond to the neural activity reflecting decision uncertainty in the brain areas such as the dACC, as evidenced in human neuroimaging studies (*22*, *23*). Both variables were calculated at the time point of decision termination, when the activity on one side of the motor module reaches the threshold. Neuroimaging studies have frequently demonstrated concurrent neural representations of both confidence and uncertainty in the brain during decision-making tasks (*22*, *23*). However, it is uncertainty, rather than confidence, that serves as the control signal for adjusting the decision-making process (*22*, *26*).

In this study, we focused on testing the model’s performance in a classical perceptual decision-making task involving judging the net direction of random dot kinematogram (RDK). The difficulty of the task was manipulated by varying the coherence of motion, representing the proportion of dots moving towards a coherent direction. For the model simulation experiments, we first reproduced the well-established relationships between decision accuracy, confidence, RTs, and the varying coherences of the RDK stimuli, and then we aimed to articulate the critical features observed in empirical studies that have been difficult to account for by the classic evidence accumulation models.

### Dissociation between confidence and uncertainty in the model

The sensory inputs in decision making usually bear noise or volatility. In the default setting, the stimulus volatility was not considered and thus the scale of input noise *β* was set as zero (equation 7 in Methods). To investigate the effect of stimulus volatility, we varied the *β* values (1, 2, 3) corresponding to different noise levels (**Fig. 3**).

**Figure 3.**
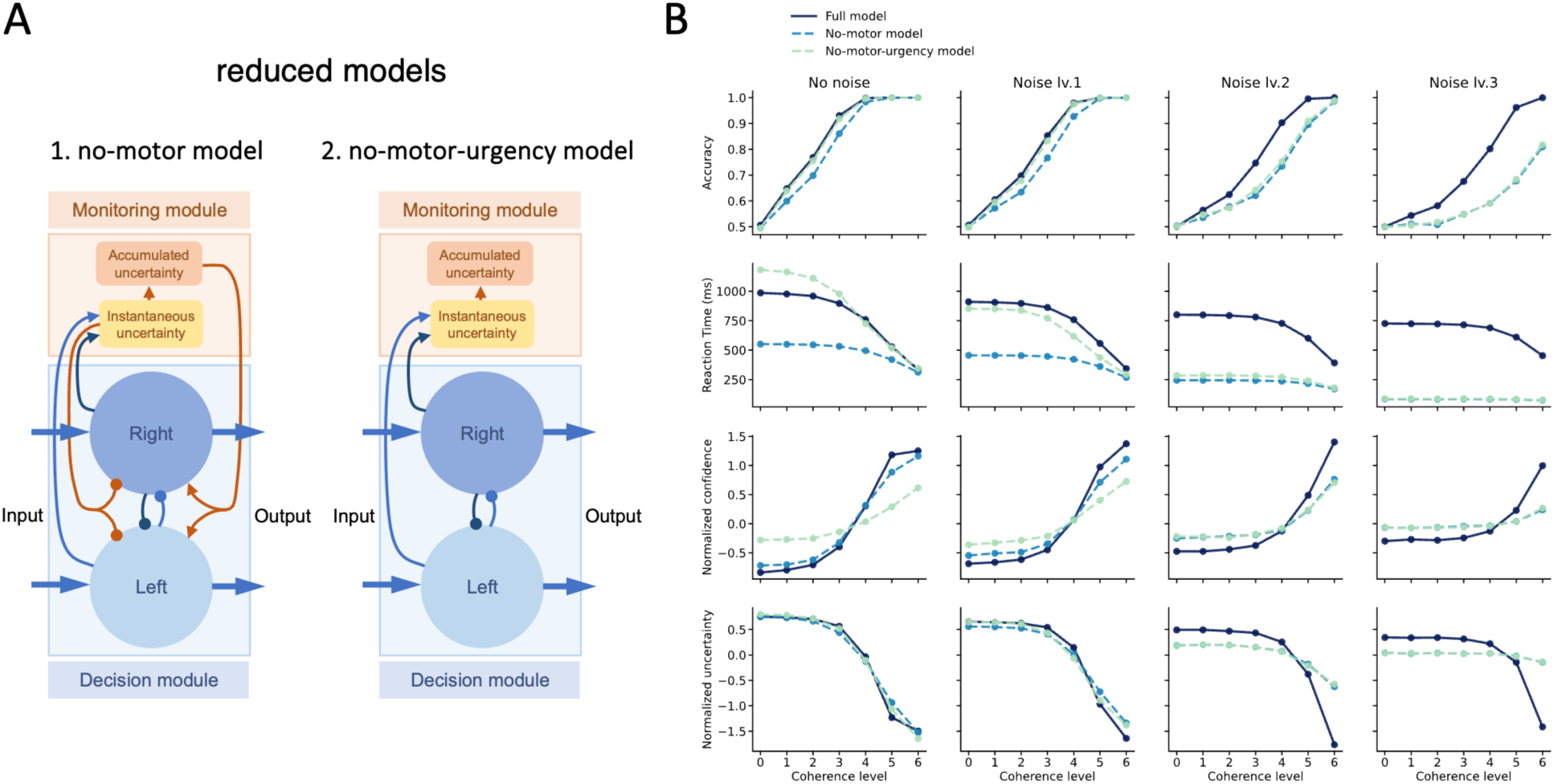
The robustness of the neural network model in faces of noise and uncertainty. (**A**) The reduced models without the motor module and further without the positive feedback from the accumulated uncertainty. Note that the strengths of mutual inhibitions between the two models were different. (**B**) The performance by the models in the faces of different levels of noise.

Although the simulations by the model successfully replicated the characteristic ‘<’-shape relationship between confidence and coherences across correct and incorrect trials (**Fig. 2B**; ref. *29*), the states of uncertainty and confidence showed distinct associations with decision accuracy and RTs at the different noise levels (**Fig. 3B**; ref. *10*). Specifically, uncertainty increased uniformly as decision accuracy decreased, regardless of the noise levels (**Fig. 4C**). By contrast, confidence had varying associations with decision accuracy at the different noise levels (**Fig. 4C**). These results imply that the indices of uncertainty more closely reflect actual decision accuracy than confidence does. Furthermore, uncertainty was orthogonally dependent on both RTs and noise levels (**Fig. 4D**), whereas confidence primarily depended on RTs with less sensitivity to the noise levels (**Fig. 4D**). The model results of uncertainty were consistent with the empirical findings that the within-trial volatility of RDK stimuli (*i.e.*, coherence variance) should lead to decreases of reaction time and increases of confidence (*10*).

**Figure 4.**
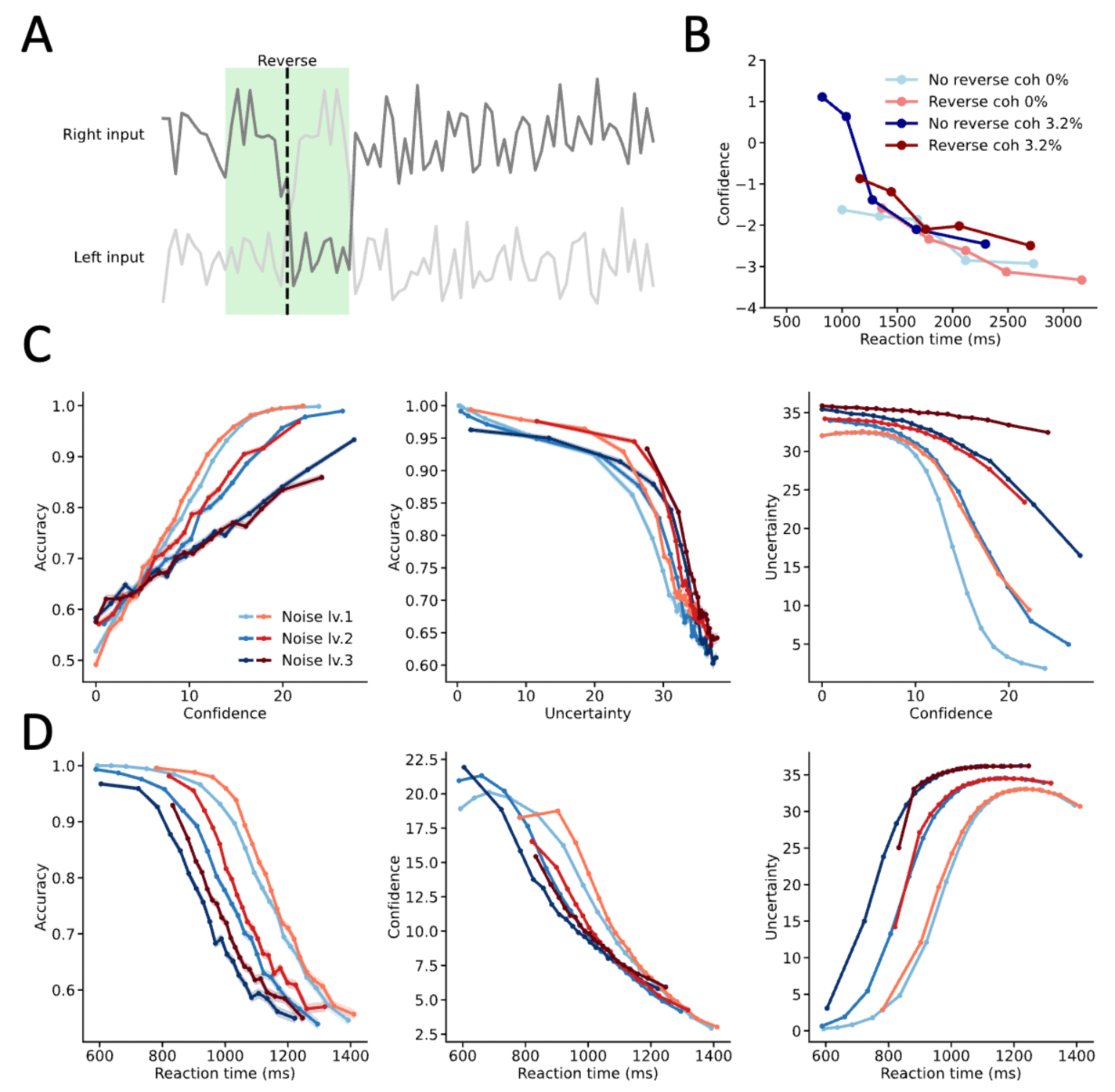
The effects of stimulus reversion simulated by the neural network model. (A) A reversal of 160-milliseconds sequence of stimulus inputs. (B) The stable inverse relationship between reaction times and confidence across different coherences and their reversions observed in the empirical study (ref. *31*). (C) The confidence-accuracy, uncertainty-accuracy and confidence-uncertainty relationships across different noise levels and their reversions. Warm colors mean reversed stimulus and cold colors mean non-reversed stimulus. (D) The reaction time-accuracy, reaction time-confidence and reaction time-uncertainty relationships across different noise levels and their reversions.

Confidence and uncertainty are internal mental states to endogenously calibrate whether the decision is correct. Hence, the measures of confidence and uncertainty should reflect both the objective decision accuracy and the subjective difficulty. A previously empirical study found that insertion of reverse stimulus pattern in the stimuli within a trial led to longer reaction time and lower confidence in human participants, while their accuracy remained almost intact (*31*). Importantly, the relationship between confidence and reaction time remained almost unchanged. Thereby, the decreased confidence should be attributed to the prolonged reaction time.

Following this empirical study, we here also inserted a reversal of 160-milliseconds sequence of stimulus inputs during the period of 200 to 360 milliseconds after stimulus onset (**Fig. 4A**). Of note, one millisecond in the model simulations corresponds to one time step. Specifically, we simulated the experiments with stimulus reversion at different noise levels. We found that the uncertainty-RT relationships, rather than confidence-RT relationships (**Fig. 4D**), remained consistent between the stimulus-reversion and non-reversion conditions (**Fig. 4B**).

Thereby, these model results illustrated that in these experiments the accumulated uncertainty might be more consistent with the empirical findings in humans, although confidence and uncertainty are conceptually two sides of the same coin. By contrast, the reduced models, lacking either the motor module or the feedback connection from the accumulated uncertainty, produced irregular uncertainty-accuracy relationships (**Fig. S5-S9**).

### Time-dependent effects of evidence on choices

The introduction of an additional stimulus pulse into the stimulus stream can also be used to investigate the time-dependent effects of new evidence on the decision-making process (*12*). Traditional decision-making models, such as the DDM, assume that new evidence arriving at each time point during the decision-making course independently contributes to the final choices (*7, 32,* but see *49*). However, empirical findings indicate that the influence of new evidence on the final choices varies over time, with a stronger impact in the early phases of decision making (*12, 33*).

In the model simulations, we followed the same procedure to investigate the time-dependent effect of stimulus influences on choices (**Fig. S1A**). The simulations were run with coherences of 0%, 1.6%, 3.2%, 6.4%, respectively. The coherence of the pulse was 12.8%, the duration of the pulse was 100 milliseconds and the direction of the pulse was either identical (positive pulse) or reverse (negative pulse) to the direction of the original stimulus. After the pulse ceased, the stimulus was altered to be a noise with 0% coherence, as a substitute for a go-cue signal immediately after the pulse offset in the empirical settings (*12*). The pulse initiated at least 50 milliseconds after the stimulus onset and when decision variable (DV) crossed a certain level of threshold that was randomly drawn from a uniform distribution in the range between 2 and 4 [*U*(2,4)] in each trial. DV was defined as the difference between the neural activities of the two competitive nodes in the motor module that were sampled per one millisecond. For those reduced models without the motor module, DV was instead defined by the difference between neural activities of the two competitive nodes in the decision module, and the pulse length was 200 milliseconds and the predefined DVs were sampled from *U*(4,6).

The pulse effects were quantified by the change of choices aligning with the original stimulation direction or DVs by comparing the conditions between the positive pulse and the negative pulse (**Fig. S1B**). However, the pulse effects might depend on both the elapsed time and the DV at the same time point. We used the same procedure as in the empirical investigation to disentangle the DV- and duration-dependent effects (**Fig. S1C**). To identify the duration effects, we sorted the data according to the DV amplitudes into ten quantiles and then subtracted the mean from the data in each quantile. We then used the residuals according different durations to quantify the pulse effects. On the other hand, we did the same procedure by removing the duration confounds to characterize the DV-dependent effects by the pulse. The analysis was completed offline and DV reflected the average in the time bin of down-sampled data. Notably, indecision trials and trials failed to initiate a pulse were not included in the analysis. The model results replicated the empirical findings that earlier pulse had a greater effect (**Fig. S1D**). The simulations were also conducted with different levels of urgency signals (*⍺*_*urg*_ =0.004, 0.006, 0.008, 0.01), and the time-related pulse effect was larger with a stronger urgency signal. Importantly, this effect was much weaker in the reduced model that lacked the motor module and the feedback from the accumulated uncertainty, but was unaffected in the other reduced models (**Fig. S5-S9**). This finding suggests that the closed-loop form of the neural circuitry is critical in generating the reduced impacts of new evidence on the final choices over time.

### Speed-accuracy trade-off

A trade-off between speed and accuracy is an important strategy deployed in decision-making (*34*, *35*). A classic perspective on speed-accuracy trade-off is the alternations of the decision threshold toward the different goals. Higher decision threshold leads to higher accuracy, but longer reaction time, and vice versa. A critical issue in the decision-making theory is how to adaptively alter the decision threshold under different objectives of speed-first or accuracy-first across contexts, trials, and even within a trial. In the current model, the most meticulous way to implement adaption of this strategy is to alter the top-down feedback from the accumulated uncertainty by adjusting the parameter of *⍺*_*urg*_ to be stronger or weaker, so that the decision threshold becomes easier or harder to be reached (**Fig. 5**). This feedback signal from the accumulated uncertainty to the motor module is thus a urgency signal (*14–16*). Hence, the generation of an urgency signal is originated from the accumulated uncertainty, a metacognitive state in monitoring the decision-making process, as illustrated by the current model. Under a relatively higher *⍺*_*urg*_, a stronger urgency signal is presented. This means that the agent might tolerate uncertainty in making a decision, and vice versa. This would lead to shorter reaction time, lower accuracy, lower confidence and higher uncertainty. There existed some alternative ways to adjust the strategy of speed-accuracy trad-off, for instance, varying the baseline of the metacognition module might lead to similar results as well.

**Figure 5.**
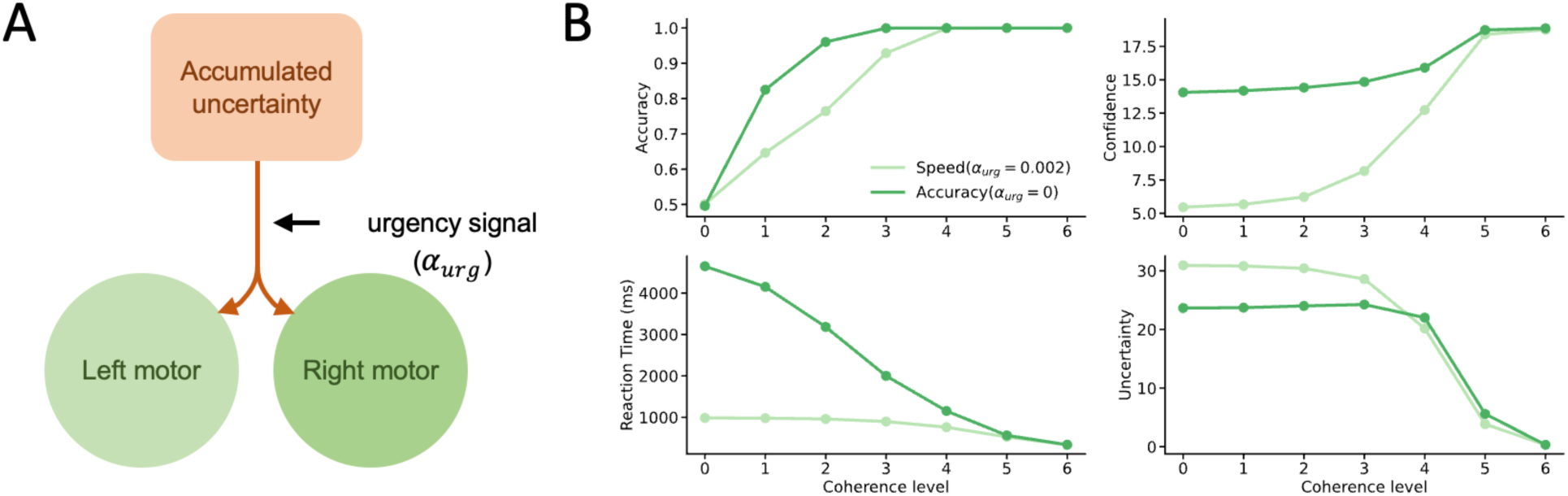
The speed-accuracy trade-off simulated by the neural network model. (**A**) The speed-accuracy strategy is simply implemented in the model by adjusting the parameter *⍺*_*urg*_ to control the urgency signal by positive feedback from the accumulated uncertainty. (**B**) Larger *⍺*_*urg*_ results in high speed, low accuracy, and high uncertainty (or low confidence).

### Attention modulations

Attention as a high-order cognitive function that may affect the decision-making process. The attentional drift-diffusion model (aDDM) was proposed to capture the modulating effect of attention on evidence accumulation (**Fig. 6B**, *36*, *37*). Briefly, it assumes that the evidence accumulation from the unattended option should be weakened relative to the attended one. As a consequence, the choices should be biased to the option that more attention has been paid on (**Fig. 6G**). In aDDM, attention is explicitly identified by gazes (*i.e.*, eye fixations; **Fig. 6F**).

**Figure 6.**
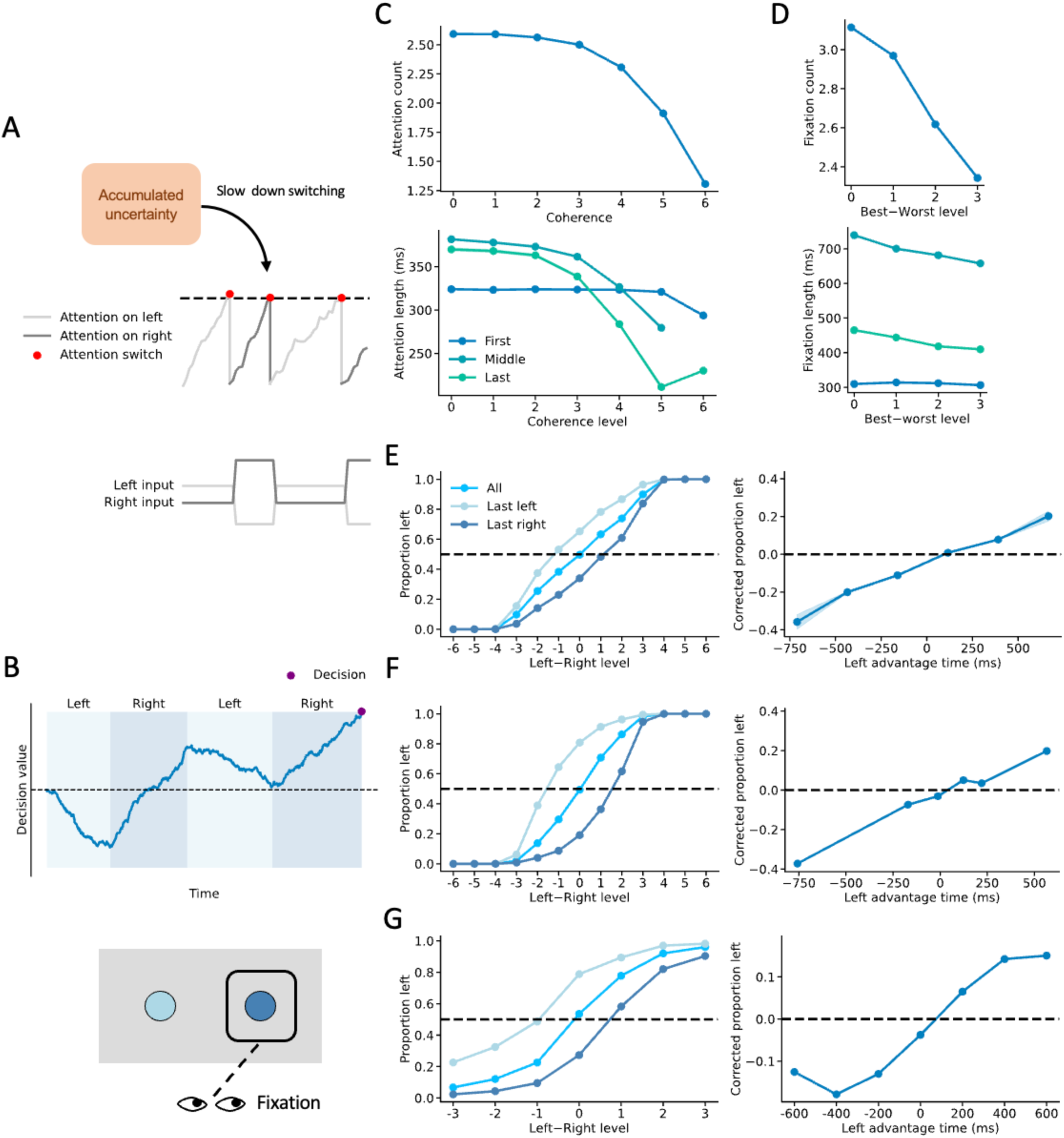
Attention modulation simulated by the neural network model. (**A**) In the model, attention to either option is assumed to be spontaneous during decision-making, while the feedback from the accumulated uncertainty might slow down switching attentions between the two alternative options. (**B**) Schema of aDDM and fixation-based attention in task. (**C**) Model results of attention counts and length. (**D**) Empirical results of fixation counts and length. (**E**) Model results of attention effect on choices. (**F**) aDDM results of attention effect on choices. (**G**) Empirical results of fixation effect on choices.

In the current model, the decision-making process is modulated by the top-down feedback from metacognition. Similarly, attention is also a cognitive control process modulated by metacognition. The orienting attention is assumed to be spontaneous and randomly switch between the two alternative options, while accumulated uncertainty is assumed to slow down the attention switching (**Fig. 6A**). The latter assumption is originated from the empirical observations that the frequency of attention switches is slowed down as the task becomes more difficult (*37*). Remarkably, the current model could predict the dynamics of attention (or eye fixations, **Fig. 6C-D**), which are taken as necessary data for the aDDM model. As a result, the current model could replicate the attention effect on choices (**Fig. 6E**).

### Change of mind

As a consequence of decision-making dynamic process, a decision can be changed during the course of the choice action until it is completed. Empirically, changes of mind (CoMs) could be explicitly detected by changes of trajectories of effector’s movements or implicitly detected by changes of trajectories of neural signals in the motor areas (*12*, *38*). CoMs often ameliorate outcomes of decision making, as changes from incorrect options to correct options (positive) occur more frequently than those from correct options to incorrect options (negative) (**Fig. S2B**).

In the current model, the detections of CoMs were based on the changes of the neural activities in the motor module (**Fig. S2A**). Specifically, an intention was defined to form when such difference was greater than 1 Hz and the dominant option remained unchanged lasting for at least 30 milliseconds, and the dominant option was referred to as the intended option. Whenever a new intention alternative to the former one was formed, a CoM was then identified. However, only the last CoM within a trial was used for the analysis. For the reduced models without the motor module, CoMs were instead identified in the neural activities of the decision module by the same approach. The criterion to detect a CoM can be changed, a similar trend has been obtained.

Prior empirical investigations found that CoMs might take place either prior or post to decision termination. The pre-decision CoMs more frequently happened in the default settings of the current model. The inclusion of the motor areas involved in the decision-making process within the current model conveniently provides a clear explanation for post-decisional CoMs (**Fig. S2**, ref. *12*, *38*), as alterations in motor neural activities directly lead to changes of choices. With a stronger urgency signal (*⍺*_*urg*_ = 0.01*nA* ⋅ *Hz*^−1^) and a delay of stimulus offset (200 milliseconds), positive-dominant post-decision CoMs could be also observed. The model results replicated the empirical observations on both the pre-decision and post-decision CoMs without new sensory inputs (**Fig. S2C**).

### Motor activity affects decision-making process

Contrary to traditional models that assume that the motor areas are not involved in decision making, the bidirectional connections between the decision module and the motor module in the current model predict that modifying neural activities in the motor module should inversely impact the neural activities in the decision module. While this inverse effect along the upstream has been demonstrated in motor planning (*39*), it had never been shown in decision making until recently. A new empirical study has reported this exact effect in a decision-making paradigm (*40*). In this study, focal unilateral inactivation of the superior colliculus (SC) through muscimol injections resulted in a requirement for stronger signals in the LIP area to terminate decisions towards the ipsilateral side (**Fig. 7A**). This translated to a higher decision boundary on the ipsilateral side, while the decision boundary on the contralateral side remained intact. Consequently, the choices were biased to option towards the intact side (**Fig. 7C**). The current model precisely reproduced these results by artificial suppression on a unilateral motor unit (**Fig. 7B**), which led to a partial reduction in the urgency signal (**Fig. 7D**). Furthermore, the current model predicted that decisions should become more uncertain (**Fig. 7E**). By contrast, the ablations on the unilateral unit within the decision module induced a shift in choices towards the contralateral side, longer RTs, decreased accuracy, and reduced confidence (**Fig. S3**). On the other hand, the ablations on the unit responsible for accumulated uncertainty, subsequently leading to a reduction in the urgency signal, thus resulted in reduced decision uncertainty, longer RTs, and improved accuracy (**Fig. S3**).

**Figure 7.**
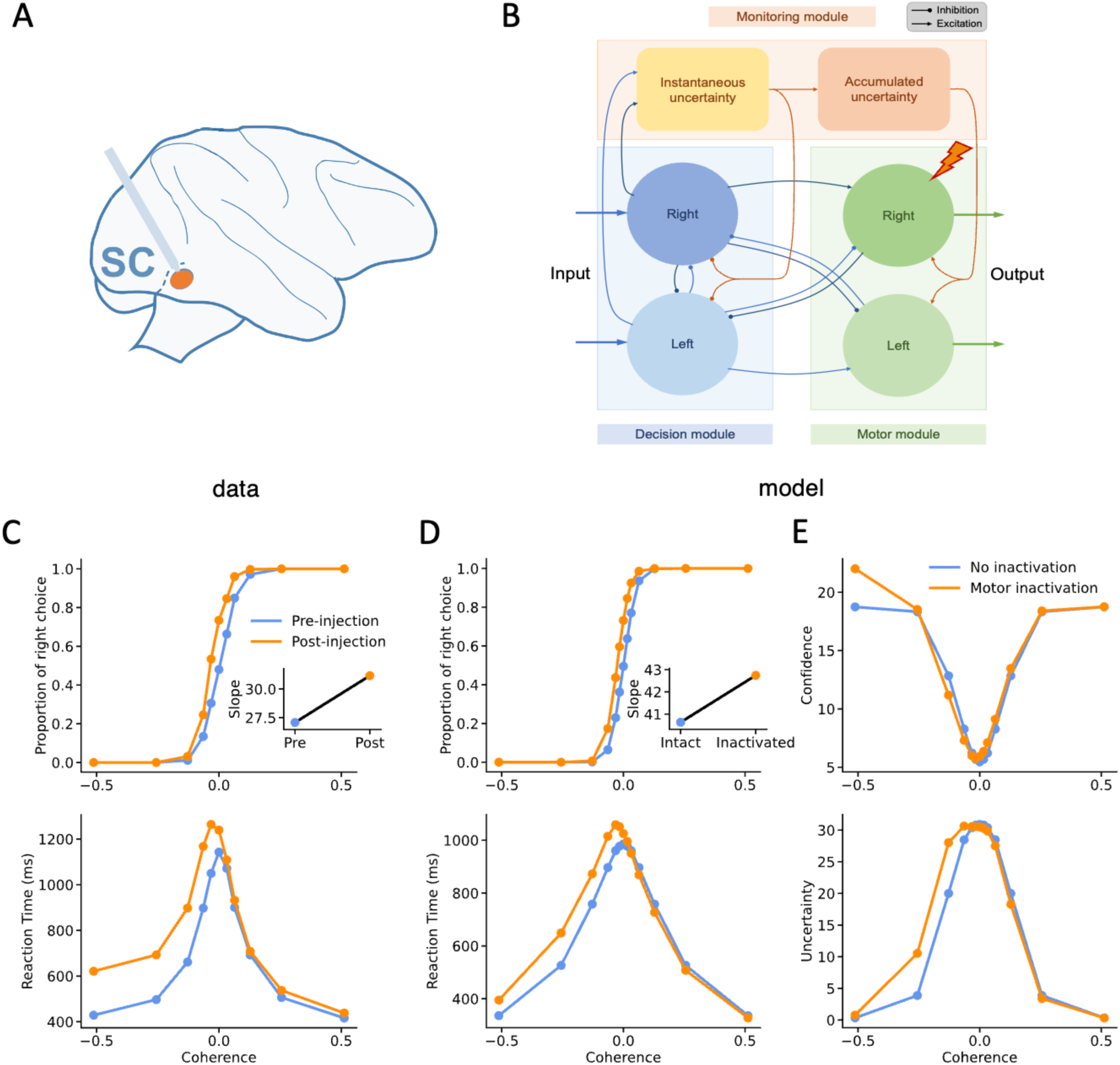
The effects by inhibitions on the unilateral motor module. (**A**) Empirical inhibitions on the unilateral superior colliculus (SC) through muscimol injections (ref. *42*). (**B**) Virtual inhibitions on the unilateral unit of the motor module in the model. (**C**) The contralateral choices and reaction times were affected in the animals’ decision-making. (**D**) A similar pattern was obtained by the unilateral inaction of the motor module in the model. (**E**) The contralateral (accumulated) uncertainty, rather than the computed confidence, was predicted by the model to be also affected.

### Decision-congruent confidence bias

A prominent phenomenon observed in decision-making revealed in recent empirical investigations is that confidence is out-weighted by the chosen option, or so-called decision-congruent confidence bias or positive evidence bias (*33*, *41*, *42*), but no bias in choices and decision accuracy (*43*). To investigate this issue by the model simulations, the coherences of stimulus inputs for the two alternative options should be independent. A basic feature for the neural system to process the sensory inputs is divisive normalization of the inputs that the neurons adaptively respond to the input signals. In other words, the neural activities are sensitive to the relative input strengths by dividing the original inputs by their summation. If the input noise all come from a constant environmental noise, the original stimulus strength *V*^j^ would be in proportion to the ratio between the input strength to the input noise 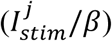. That is, 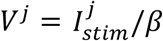, we then defined *ΔV* = *V*^*chosen*^ − *V*^*unchosen*^ and *ΣV* = *V*^*chosen*^ + *V*^*unchosen*^. We used |*ΔV*|, *ΣV* and reaction time to fit decision accuracy, and used *ΔV*, *ΣV* and reaction time to fit confidence and uncertainty in the simulations by the models. Thereby, a significant weight on the *ΣV* variable then indicates a confidence bias, when the weight on the *ΔV* variable is also significant.

Among the simulated data with noise strength *β* = 1, 2, 3, we uniformly draw a batch of 300 samples from trials with non-zero coherence. For reduced models without the motor module, we used the data with weaker noise where *β* = 0.5, 1, 1.5. The model confidence and uncertainty were transformed into [0,1] by min-max normalization in each batch. Binomial generalized linear model with logit link was used to fit the data.

We repeated this process 100 times. The p-value was obtained based on percentile rank of 0 in these distributions. We found that in the full model larger *ΣV* led to lower uncertainty even though its effect on decision accuracy was not significant (**Fig. S4**). By contrast, the effect of *ΣV* on model confidence was not significant. Such a significant effect of *ΣV* in modulating uncertainty equals to a larger weight of *V*^*chosen*^ on uncertainty in comparison to *V*^*unchosen*^. These results replicated the empirical observations (*33*, *41–43*). Notably, our model results show that the accumulated uncertainty, rather than the calculated confidence, reliably replicated such a selective bias (**Fig. S4**), even in the reduced models (**Fig. S5-S9**).

## Discussions

Taken together, in the current study we have demonstrated that the proposed model, which synergistically integrates the motor module, the metacognition module, and the traditional decision module, effectively captures the dynamic features of decision making and metacognition, particularly in the presence of noise and uncertainty. Not only does the model replicate classical findings in decision making and metacognition, but it also successfully predicts some new critical empirical results that traditional models may fail to explain. For instance, the adaptive speed-accuracy tradeoff (**Fig. 5**), motor activity influencing decision-making process (**Fig. 7**), pre-decisional and post-decisional change of mind (**Fig. S2**). Remarkably, although uncertainty and confidence are two sides of the same coin, the explicit representation of uncertainty in the metacognition module of the model, more accurately and stably than the estimated confidence by ‘the balance of evidence’ (*11*, *29*, *30*), reflect decision accuracy under different levels of uncertainty (**Fig. 3** and **Fig. 4**). These remarkable replications of empirical findings can be attributed to the underlying neural network architecture embedded in the model. Importantly, our results demonstrate that these components in the model are indispensable for replicating these decision-making behaviors.

A significant advance in the decision-making theory presented by this model is the establishment of an inherently reciprocal relationship between decision making and metacognition. While previous studies focused on how decision uncertainty or confidence is generated from the decision-making process (*9–11*), the current model delineates how metacognition monitors and regulates the decision-making process (*26*, *27*, *44*). It illustrates that decision uncertainty is not merely a by-product of decision making but, in fact, plays crucial roles in shaping the decision-making process. The cognitive control theory has also suggested to modulate the decision-making process in a self-regulating manner (*45–47*). However, the top-down modulation is sequentially induced by decision making. Thereby, it emphasizes an after-effect control on the sequential task performance, in particular, multiple tasks in parallel, rather than the dynamic control within the currently ongoing decision-making process, as proposed in the current model. A recent study has also proposed a neural network model similarly consisting of the decision, motor and metacognition modules, in which the uncertainty unit excitably feeds back to the decision module, without feedbacks toward the motor module and feedbacks from the motor module to the decision module (*24*). It might correspond a simplified model where the decision modules and motor modules in our model is collapsed together. One unique feature of this model distinguishing from our current model is that the motor module is passively activated by receiving inputs from the decision module, but lacks interactions with the decision module and the metacognition module. One drawback of such a neural network architecture consisting of one direct loop between the decision module and the metacognition module is that the decisions under uncertainty are speedily determined without deliberation. As a consequence, the accuracy becomes much lower under higher uncertainty (Fig. 2 in ref. 24). Namely, the decisions are suboptimal. By contrast, the current model through dual regulations from the metacognition module separately on the decision module and the motor module dynamically balances deliberation and urgency for optimal decision making.

The inclusion of the motor module as an integral part of the decision-making process in the current model, particularly by decoupling the dynamics of the two dual modules, is critic to enhance the resilience of decision making to noise and facilitates adaptability in changing circumstances. The involvement of motor areas in decision making has been repeatedly showed by empirical (*12*, *13*, *40*) and modelling (*48*) studies.

Although the urgency signal has been recognized to be crucial for optimal decision making (*5*, *16*), its neural origin remains unclear (*49*). The current model provides an inherent mechanism in generation of an urgency signal through the accumulation of momentary decision uncertainty, which arises from the neuronal activities associated with evidence accumulation in the decision module. This urgency signal then executes on the motor module to expedite the choice commitments (*50*), in particular, effectively at the late phase of decision making. The dynamic modulation from the metacognition module to the motor module thus leads to adaptive adjustment of the speed-accuracy tradeoff strategy. Hence, optimality in decision making is an intrinsic property of the decision-making neural circuity as illustrated in the current model. Notably, in our model, the urgency signal generated by the accumulated uncertainty node effectively reduces the threshold for terminating decision making as deliberation lasts toward the late phase. Such a modulation on the decision-making process by the urgency signal is different from that in the other models that the urgency signal serves as global gain modulation on the two-alternative evidence accumulation (the urgency signal multiplies the momentary accumulated evidence, ref. *15*, *18, 49*), corresponding to increases of drift rates along the course of evidence accumulation in the DDM model. Hence, the latter manner of modulation by the urgency signal is exactly an adjustment of unselective attention during decision making. Although this modulation should occur during decision making, it still has limited capacity to accelerate evidence accumulation to reach the stationary threshold when the evidence is too weak, for instance, at 0% coherence in the RDK task. Accordingly, the negative modulation on the decision module from the instantaneous uncertainty node in our model might have an equivalent effect as a global gain by timely suppressing noise from the two competitive sensory inputs, in particular, during the early phase of decision making.

On the other hand, metacognitive signals may modulate gating of the inputs through selective attention during the decision-making process. Higher uncertainty signals induce more selective attention on the inputs but in a stochastic manner during the decision-making process. Although this modulation circuit is not implemented in the current model, we may use the accumulated uncertainty signals to produce the latent attention dynamics and their effects on choices, similarly as the aDDM model predicts (*36*, *37*, **Fig. 6**). Importantly, our model may even predict the latent states of attention, for instance, indicated by the gaze positions (*36*, *37*).

In summary, the current theoretical study demonstrates that decision making is not solely a process of bottom-up evidence accumulation, but rather an adaptive control system that incorporates a closed-loop neural network architecture of decision-making and metacognition. The proposed model provides a plausible neural implementation that can account for many critical empirical findings in humans and animals, and importantly, make some predictions about the interacting dynamics of decision-making and metacognition that can be testable in the future neural and behavioral studies. Remarkably, this model highlights a deliberative decision-making process under uncertainty in pursuit of optimality (*16*, *28*), which may differ from ballistic or fast decision-making processes observed in native and over-trained tasks (*51–53*). The current model provides insights on the neural circuit mechanisms of decision making and metacognition, shedding light on the underlying neural dynamics, and potentially identifying the pathological and psychiatric sources of irrational decision-making behaviors in patients with mental disorders.

## Materials and Methods

### Intuitive link between instantaneous uncertainty and entropy

In our model, we use the uncertainty module to track the amount of contradictory information. This is consistent with existing hypothesis about the functions of some uncertainty-related brain areas, which we know little about their neuro-computational mechanism. As a result, it would be beneficial to have an intuition of why the framework could lead to such functions in our model.

Let us consider information entropy *H* = −∫ *p*log*p* and its variant *H*^∗^ = *H* + 1 = ∫ (*p* − *p*log*p*), where *f*(*p*) : = *p* − *p*log*p* monotonically increases as *p* ∈ [0,1]. Besides, note that *f*(*p*) is concave as *f*″(*p*) = −1/*p* < 0, i.e., *f*(*p*) increases slower with larger *p*. Such monotonicity and saturation characteristic could also be seen in the excitatory inputs from decision modules to the instantaneous uncertainty module in our model. These inputs increase with decision module activities, and increases slower with higher activities due to saturation of connections. From this point of view, we could link the instantaneous uncertainty in our model to *H*^∗^, and treat it as a proxy of entropy. Thus, the idea that the instantaneous uncertainty module tried to minimize the activities of itself corresponds to minimizing entropy.

### The neural-network model

Following the prior research (*6*, *24*, *25*, *54*), our model is built in a reduced form of neural network described by a mean-field approximation of populational neuronal activities with slow dynamics of NMDA gating for each node of the decision, motor and metacognition modules in the model (*6*). That is,

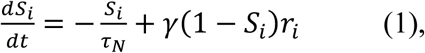

where *r*_*i*_ denotes the firing rate of node *i* and *S*_*i*_ denotes the NMDA-gating variable related to synaptic connections. Further, the dynamics of *r*_*i*_ depends on the current input *I*_*i*_ with an input-output function *ϕ*:

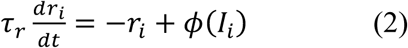

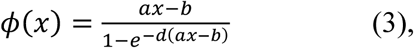

where *I*_*i*_ can be further divided into components of constant background 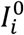, input 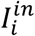, recurrent connection 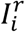, and noise 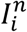:

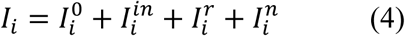

In this model, all other nodes share the same baseline input strength *I*^0^ with an exception that the instantaneous uncertainty node has an additional baseline inhibition,

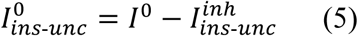

and

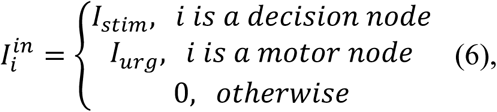

where the inputs to the nodes in the decision module are the stimulus coherences *∈* and are modulated by independent noises as follows.

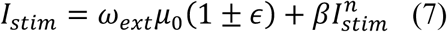

The urgency signal inputs to the nodes of the motor module are assumed to be proportional to the firing rate of the accumulated uncertainty node,

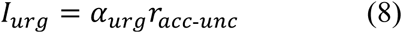

and

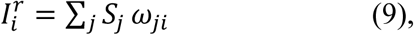

Where *⍺*_*urg*_ and *ω*_j*i*_ denotes the modulation strength from the accumulated uncertainty node and the strengths of inter-node or within-node recurrent connections from node *j* to node *i*.

Specifically, a positive value indicates an excitatory connection and a negative value indicates an inhibitory connection.

All nodes have the same noise as an Ornstein-Uhlenbeck process:

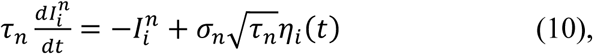

where *η*_*i*_ is Gaussian white noise with zero mean and unit variance.

For sake of simplicity, the nodes in all modules of the current model share the same formulas and parameter values. Notably, we use similar but simpler formulas with *τ*_*acc*_ ≫ *τ*_*r*_ for the dynamics of accumulated uncertainty node.

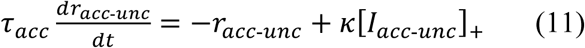

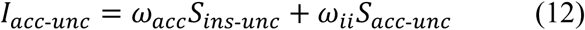

Based on these formulas, the trial-by-trial decision-making processes were simulated. When the firing rate of either node in the motor module crossed the predefined threshold, a decision was made and the stimulus presentation ceased. The choice then corresponded to the side of nodes that crossed the threshold and its reaction time was the elapsed time since the stimulus onset. Confidence was calculated as the difference between the firing rates of decision nodes (‘balance of evidence’, *11*, *29*, *30*), while uncertainty was defined as the activity strength of the accumulated uncertainty node at the time point of decision termination.

### Implementation details

The parameters of the local neural circuit were adapted from the prior studies (*6*, *24*), while additional parameters specific to the new components of the model were customized to capture the fundamental properties of decision making and metacognition (**Table S1-S5**). In each trial, the stimulus will be on after 300 milliseconds of preparing time and this was not included in the calculation of reaction time. The decision threshold was chosen to be 30 Hz and the maximum trial length was 3,000 milliseconds. If a decision was incomplete within this duration since the stimulus onset, then this trial was considered as a void trial and the data of this trial was excluded except for the accuracy calculation. However, due to the emergence of an urgency signal, no void trials occurred in the default settings. While investigating the speed-accuracy trade-off strategies during decision making, the maximum trial length was extended to 15,000 milliseconds, in order to obtain comparable trials in the accuracy-first instances. In each simulation setting with a certain level of stimulus coherence for each model, we simulated 10,000 trials. The confidence and uncertainty levels were calculated by averaging the dynamic data in a window of 10 milliseconds prior to the time point of terminating the decision-making process, while other data analyses were completed offline based on down-sampled data by time bins of 30 milliseconds. The stimulus coherences were 0%, 1.6%, 3.2%, 6.4%, 12.8%, 25.6%, 51.2%, respectively, and one time step updated for the decision-making dynamics was set to be 1 millisecond. Besides, by definition, our model’s uncertainty is a positive value. However, due to task difficulty and noise, model’s confidence value could be negative. To make it more reasonable, we transformed the negative confidence values to be zero.

### Attention modulation details

We added attention modulating of input into our model and further assumed that accumulated uncertainty would slow down the attention switching. The latter assumption came from the relationship between task difficulty and attention switches observed in experiments. Thus, unlike aDDM, our model could generate attention data rather than depends on existing experiment data. Specifically, we assumed that the gain of evidence accumulation of the attended option increased by a factor of 1.2, while the attention switches were generated through a random process as follows,

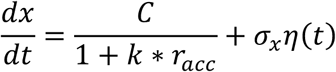

where 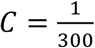, *k* = 0.1, *σ* = 0.01 and *η* is Gaussian white noise with zero mean and unit variance. The initial value of *x* is zero, and as soon as *x* reach the threshold (*b* = 1), an attention switch occurs and *x* returns to the initial point. The option to be initially attended is also chosen randomly with a uniform probability. The model results qualitatively replicated the empirical results (*36*, *37*) and aligned well with the aDDM predictions.

For a fair comparison, the attention data used for the aDDM model was generated by our model. In other words, in each trial of simulation by our model, we ran a corresponding simulation using the aDDM model. In each trial, the coherence *ε* was matched and aDDM received inputs *I*_j_ = 1 ± *∈*, with *x*_0_ = 0, the dynamics of aDDM evolution was formulated by

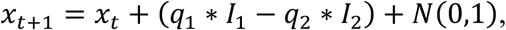

where *q*_j_ took the value of 1 if option *j* was attended and 0.7 otherwise. We assumed each time step in aDDM also corresponded to a time step in our model. Thus, at each time step, the attended option in aDDM could be determined by corresponding attention data of our model. Whenever |*x*_*t*_| exceeded a threshold of 100, aDDM made an independent choice. If aDDM did not reach the threshold within the model reaction time, additional attention lengths were sampled from distribution of the middle attention length generated by model simulation under the same stimulus coherence.

### Module inactivation details

We tried to add constant inhibition to other modules to see their effect in the decision-making process. To be detailed, the inhibition strength was 0.002 *nA* for the decision module, 0.01 *nA* for the motor module and 10 *nA* for the accumulated uncertainty node in the full model with the default settings of parameters as described above.

### Reduced models

We also interrogated the following five reduced models by ablations of some components or modules of the full model: 1. ‘no-motor-feedback’ model (**Fig. S5**); 2. ‘no-instantaneous uncertainty-feedback’ model (**Fig. S6**); 3. ‘no-motor’ model (**Fig. S7**); 4. ‘no-urgency’ model (**Fig. S8**); 5. ‘no-motor-urgency’ model (**Fig. S9**). The motor module was not included in the ‘no-motor’ model and the ‘no-motor-urgency’ model. In these models, A decision threshold of 20 *Hz* was set at the decision module and the strength of mutual inhibition between the two nodes in the decision module was set to 0.0497 *nA*, as same as in the classical decision-making model (*6*, *24*). Notably, the top-down feedback from the metacognition module was reserved in the ‘no-motor’ model, but were not included in the ‘no-urgency’ model and in the ‘no-motor-urgency’ model. In the ‘no-instantaneous uncertainty-feedback’ model, the inhibitory feedback from instantaneous uncertainty to the nodes of the decision module was removed, and the strength of mutual inhibition was set to 0.0497 *nA*. In the ‘no-motor-feedback’ model, only the feedback from the motor module to the decision module was removed. For models with strong mutual inhibition between decision modules, the strength of the additional baseline inhibition for the instantaneous uncertainty module was changed to 0.1 *nA*.

## Author contributions

Conceptualization: WL, XW; Methodology: WL, XW; Investigation: WL, XW; Visualization: WL, XW; Funding acquisition: XW; Project administration: XW; Supervision: XW; Writing – original draft: WL, XW; Writing – review & editing: XW.

## Data and materials availability

The codes used in the current study can be reached through https://github.com/Lu-WW/closed-loop-model.

---

**Table S1.** Full model parameter Values.

| Parameter | Value | Remarks |
| --- | --- | --- |
| $a$ | $270 \text{ Hz} \cdot \text{nA}^{-1}$ | refer to previous models (6, 24) |
| $b$ | $108 \text{ Hz}$ | refer to previous models (6, 24) |
| $d$ | $0.154 \text{ s}$ | refer to previous models (6, 24) |
| $\kappa$ | $1 \text{ Hz} \cdot \text{nA}^{-1}$ | manually chosen |
| $\mu_0$ | $30 \text{ Hz}$ | refer to previous models (6, 24) |
| $\omega_{ext}$ | $0.00052 \text{ nA} \cdot \text{Hz}^{-1}$ | refer to previous models (6, 24) |
| $I^0$ | $0.3255 \text{ nA}$ | refer to previous models (6, 24) |
| $I_{ins-unc}^{inh}$ | $0.2 \text{ nA}$ | manually chosen; |
| $\tau_N$ | $100 \text{ ms}$ | refer to previous models (6, 24) |
| $\gamma$ | $0.641$ | refer to previous models (6, 24) |
| $\tau_{noise}$ | $2 \text{ ms}$ | refer to previous models (6, 24) |
| $\tau_r$ | $2 \text{ ms}$ | refer to previous models (25) |
| $\omega_{ii}$ | $0.24 \text{ nA}$ | refer to previous models (24) |
| $\sigma_{noise}$ | $0.02 \text{ nA}$ | manually chosen |
| $\tau_{acc}$ | $100 \text{ ms}$ | manually chosen |
| $\omega_{acc}$ | $50 \text{ nA}$ | manually chosen |
| $\alpha_{urg}$ | $0.002 \text{ nA} \cdot \text{Hz}^{-1}$ | manually chosen |
| $\beta$ | $0$ | no noise as default |
| $I_{ins-unc}^0$ | $0.1255 \text{ nA}$ | $I^0 - I_{ins-unc}^{inh}$ |

| From | To | $\omega_{from,to}$ |
| --- | --- | --- |
| decision | opposite decision | $-0.005 \text{ nA}$ |
| decision | opposite motor | $-0.3 \text{ nA}$ |
| motor | opposite decision | $-0.02 \text{ nA}$ |
| decision | corresponding motor | $0.2 \text{ nA}$ |
| decision | instantaneous uncertainty | $0.25 \text{ nA}$ |
| instantaneous uncertainty | decision | $-0.02 \text{ nA}$ |

**Table S2.** Modified parameters for the ‘no-instantaneous uncertainty-feedback’ model.

| Parameter | Value | Remarks |
| --- | --- | --- |
| $I_{ins-unc}^{inh}$ | 0.1 nA | altered baseline |
| From | To | $\omega_{from,to}$ |
| decision | opposite decision | -0.0497 nA |
| instantaneous uncertainty | decision | 0 nA |

**Table S3.**
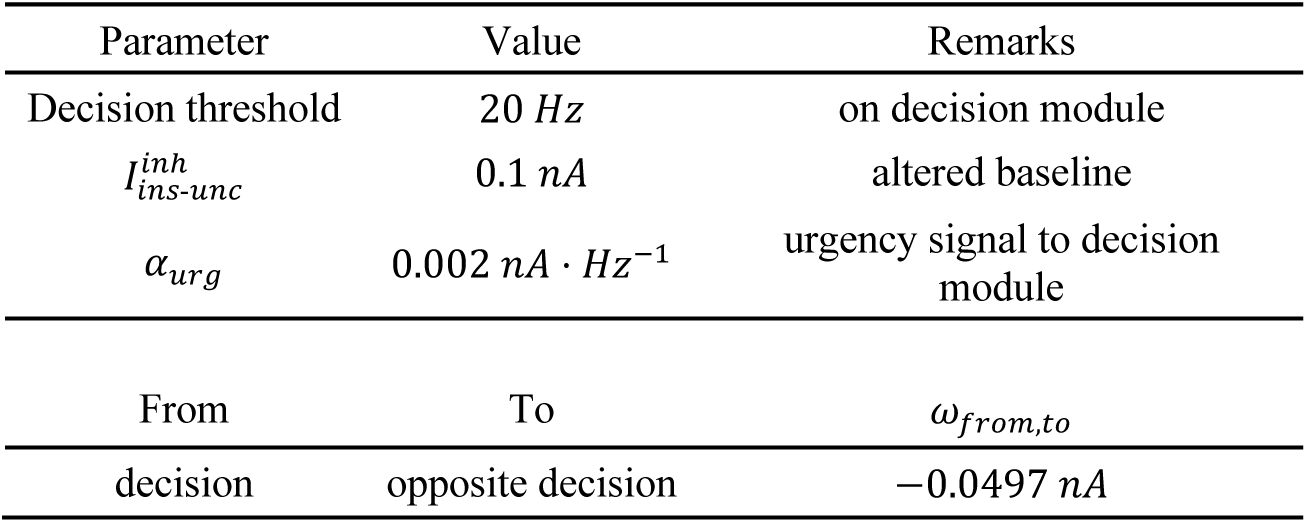
Modified parameters for the ‘no-motor’ model.

**Table S4.**
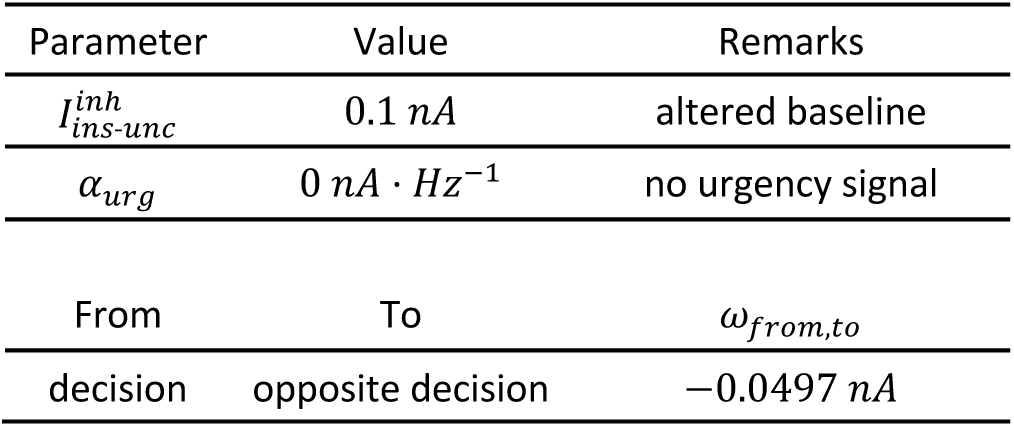
Modified parameters for the ‘no-urgency’ model.

**Table S5.**
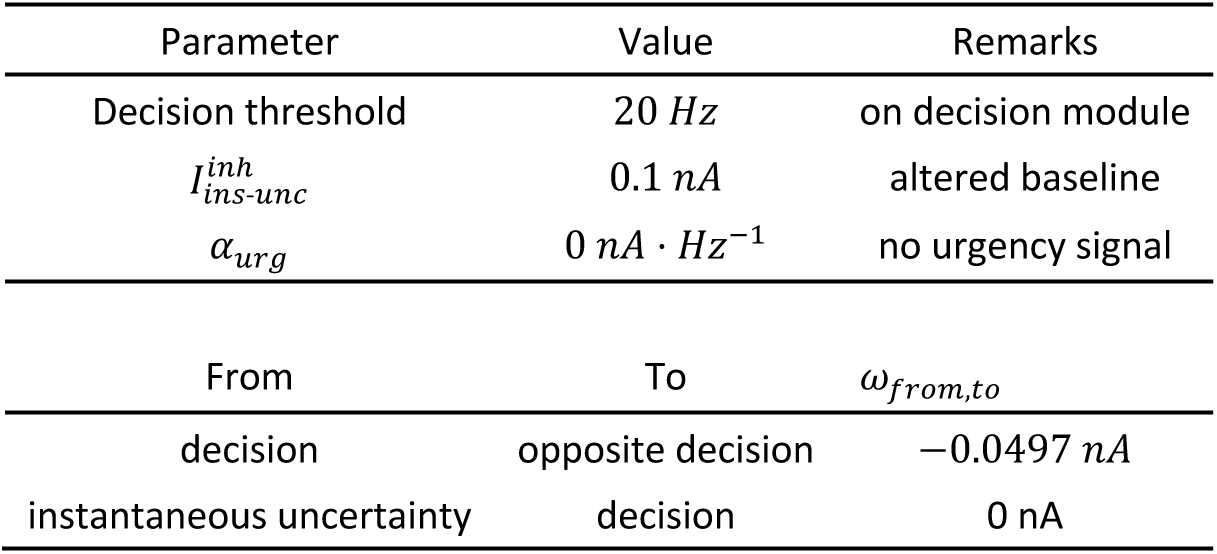
Modified parameters for the ‘no-motor-urgency’ model.

| Parameter | Value | Remarks |
| --- | --- | --- |
| Decision threshold | $20 \text{ Hz}$ | on decision module |
| $I_{ins-unc}^{inh}$ | $0.1 \text{ nA}$ | altered baseline |
| $\alpha_{urg}$ | $0 \text{ nA} \cdot \text{Hz}^{-1}$ | no urgency signal |
| From | To | $\omega_{from,to}$ |
| decision | opposite decision | $-0.0497 \text{ nA}$ |
| instantaneous uncertainty | decision | $0 \text{ nA}$ |

**Figure S1.**
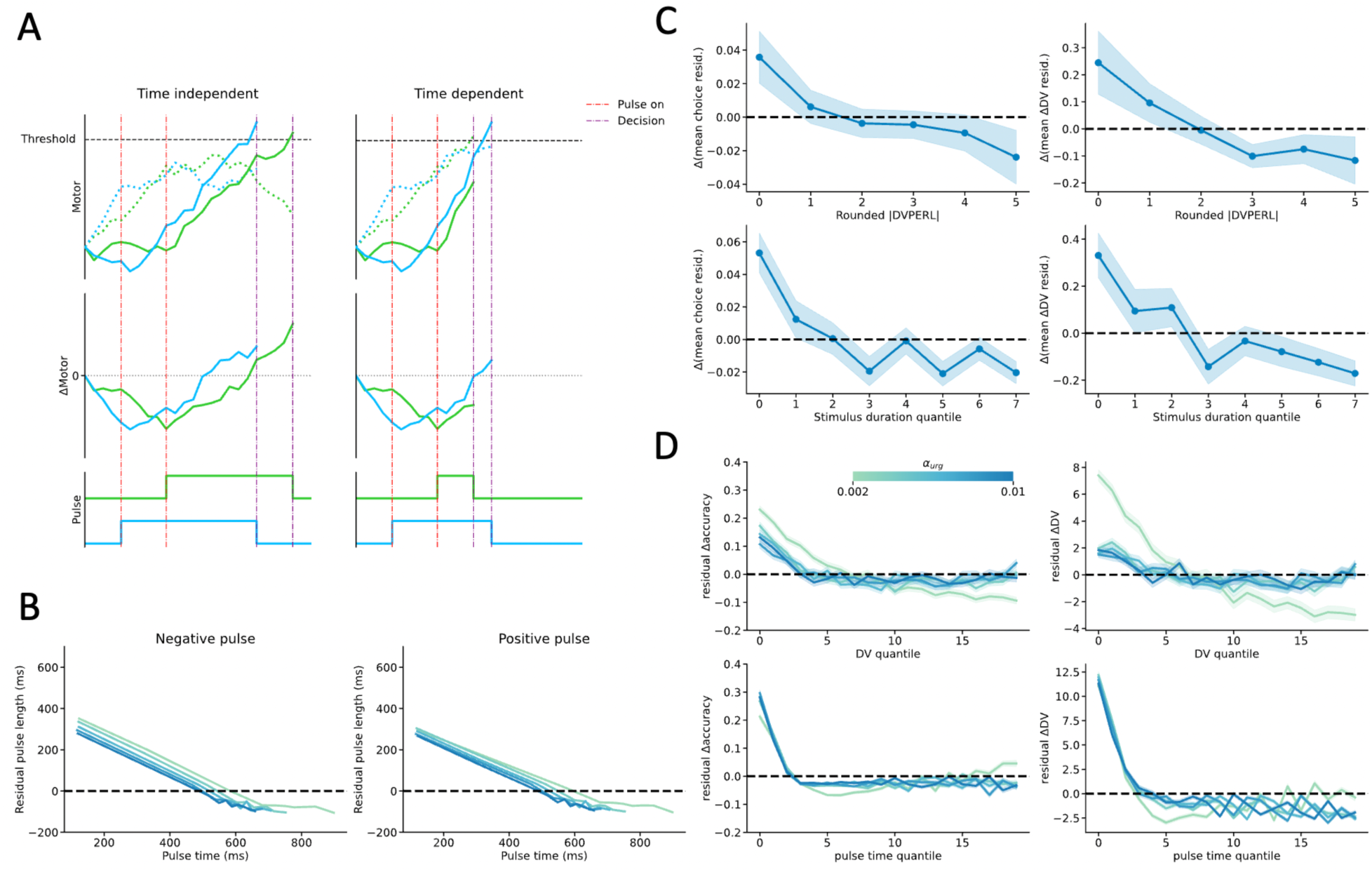
The effects of stimulus pulse simulated by the neural network model. (**A**) The effective duration of a certain stimulus pulse used to test its effects on the choices and decision variables (DVs) is different. (**B**) The effective duration was dependent on the onset of stimulus pulse presentation. The earlier the onset, the longer the effective duration. (**C**) The effects of stimulus pulse on the choices (left column) and the DVs (right column) depended on the accumulated DVs (upper) and the onset of stimulus pulse (bottom) after controlling the confounding effects in the empirical study (ref. *12*). (**D**) The effects of stimulus pulse on the choices (left column) and the DVs (right column) depended on the accumulated DVs (upper) and the onset of stimulus pulse (bottom) after controlling the confounding effects simulated by the model.

**Figure S2.**
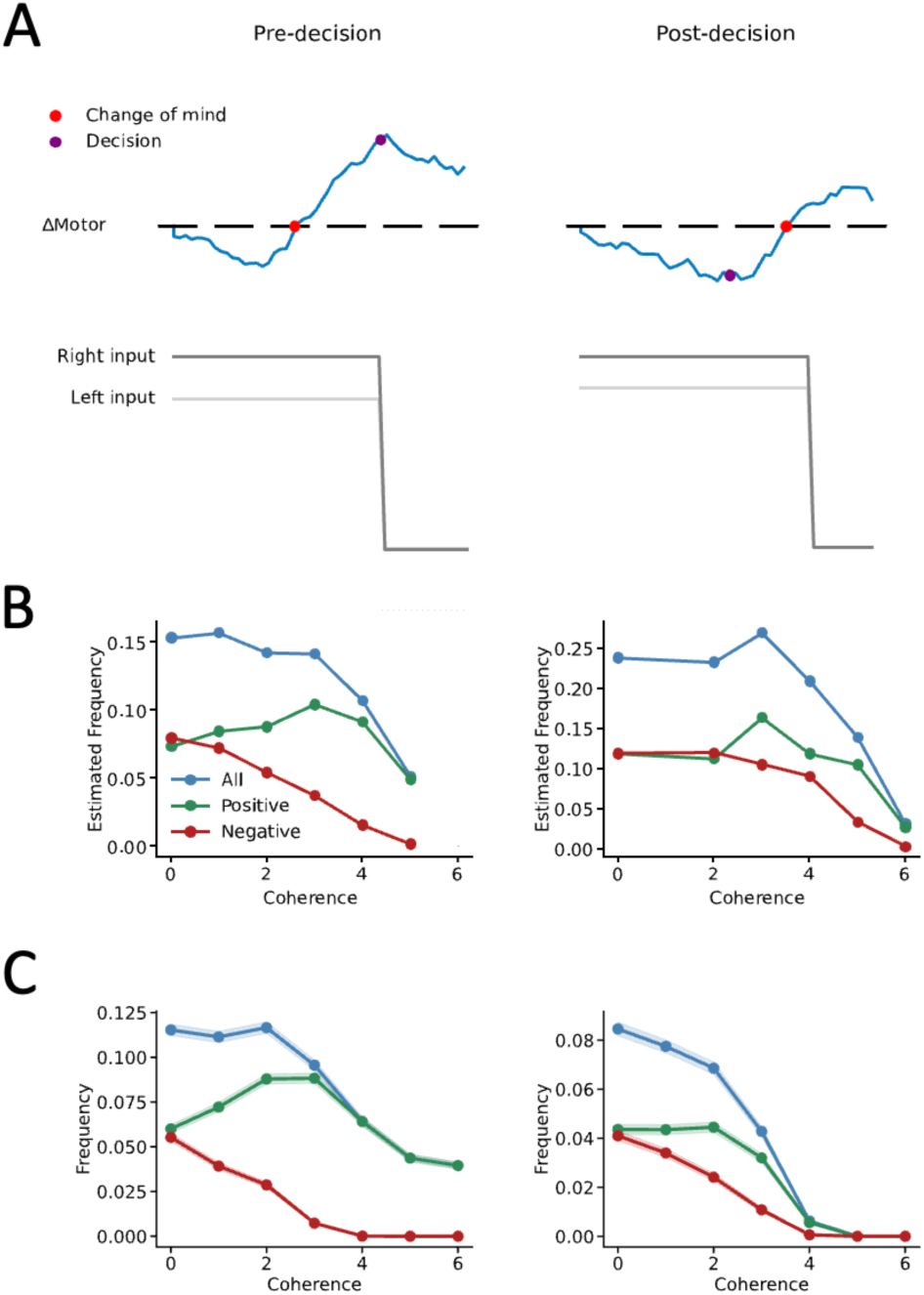
Change of mind (CoM) simulated by the neural network model. (**A**) The definition of pre-decisional and post-decisional CoM when the changes of decision variable cross the threshold. (**B**) The pre-decisional (ref. *12*) and post-decisional (ref. *38*) CoMs observed in the empirical study. (**C**) The pre-decisional and post-decisional CoMs simulated by the model.

**Figure S3.**
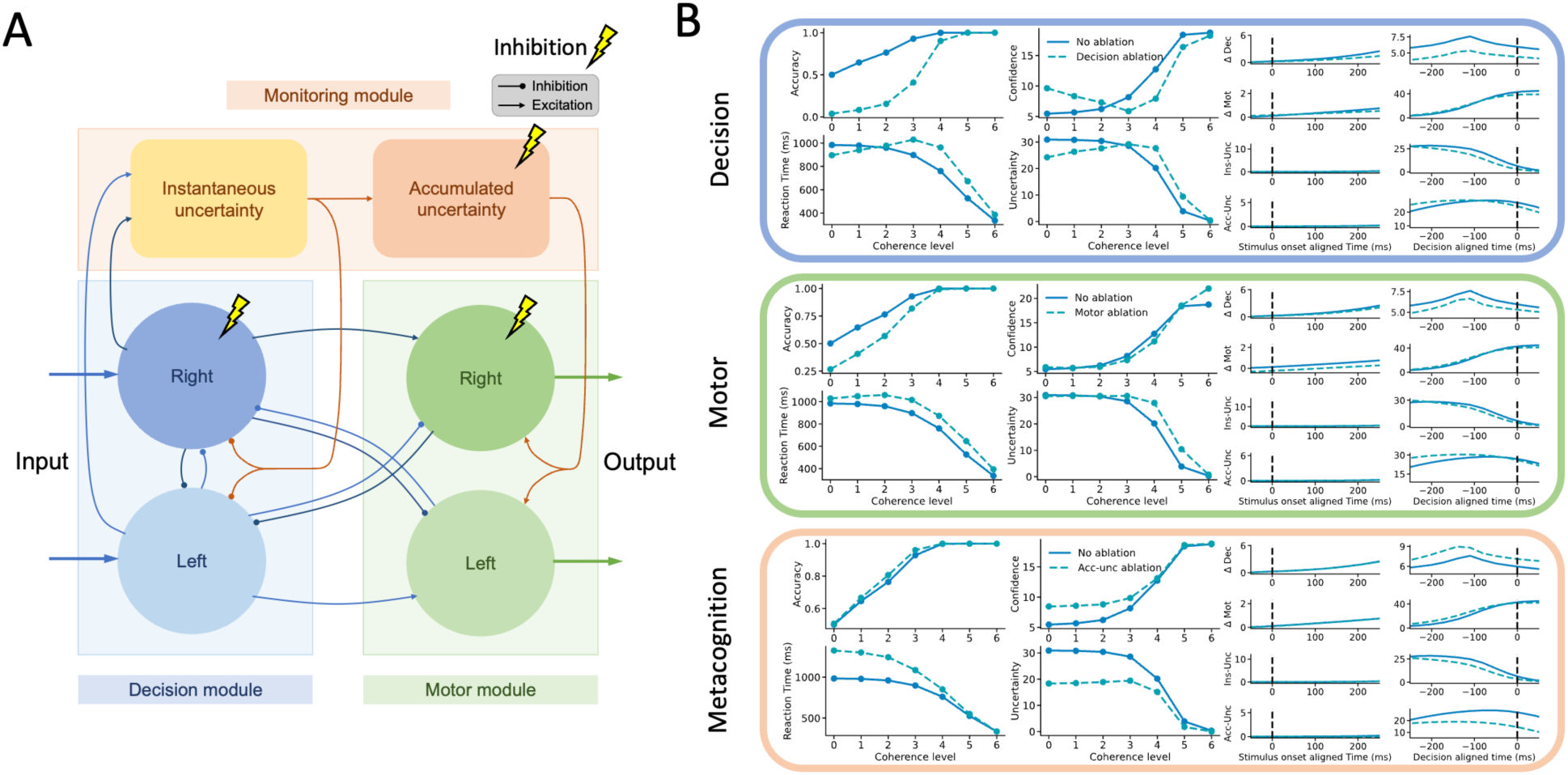
Module inactivation in the neural network model. (**A**) Independent inactivation of the unit of the decision module, the motor module and the accumulated uncertainty by adding a constant negative current for a duration to the target unit. (**B**) The neural and behavioral effects by each module inactivation.

**Figure S4.**
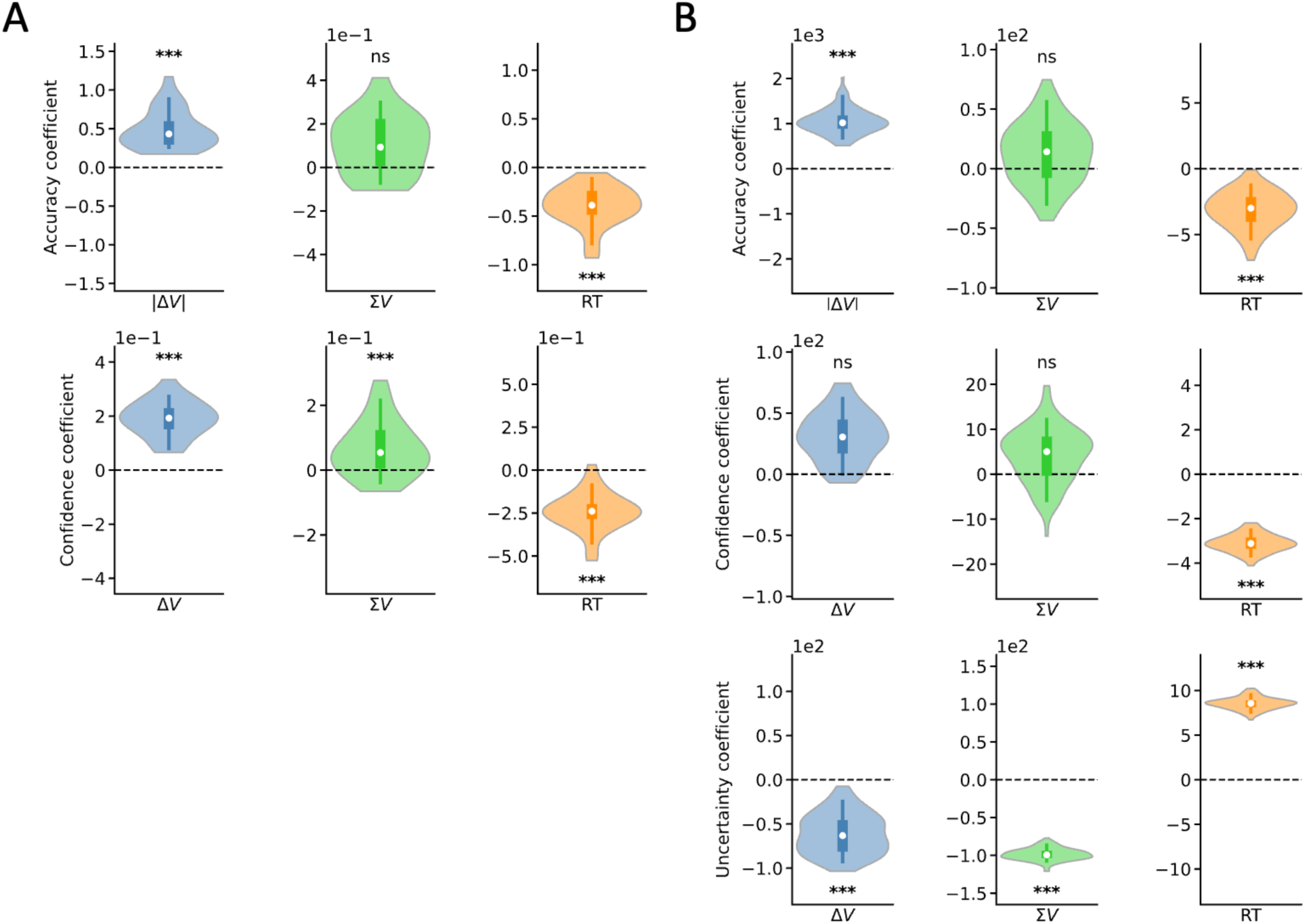
Decision-congruent confidence bias simulated by the neural network model. (**A**) The decision accuracy is dependent on the value difference (*ΔV*), but not the value sum (*ΣV*), between the two alternative options; by contrast, the confidence in believing that the choice is correct was dependent on both *ΔV* and *ΣV*, in our empirical study. (**B**) The model replicated the selective bias in uncertainty, but not in confidence.

**Figure S5.**
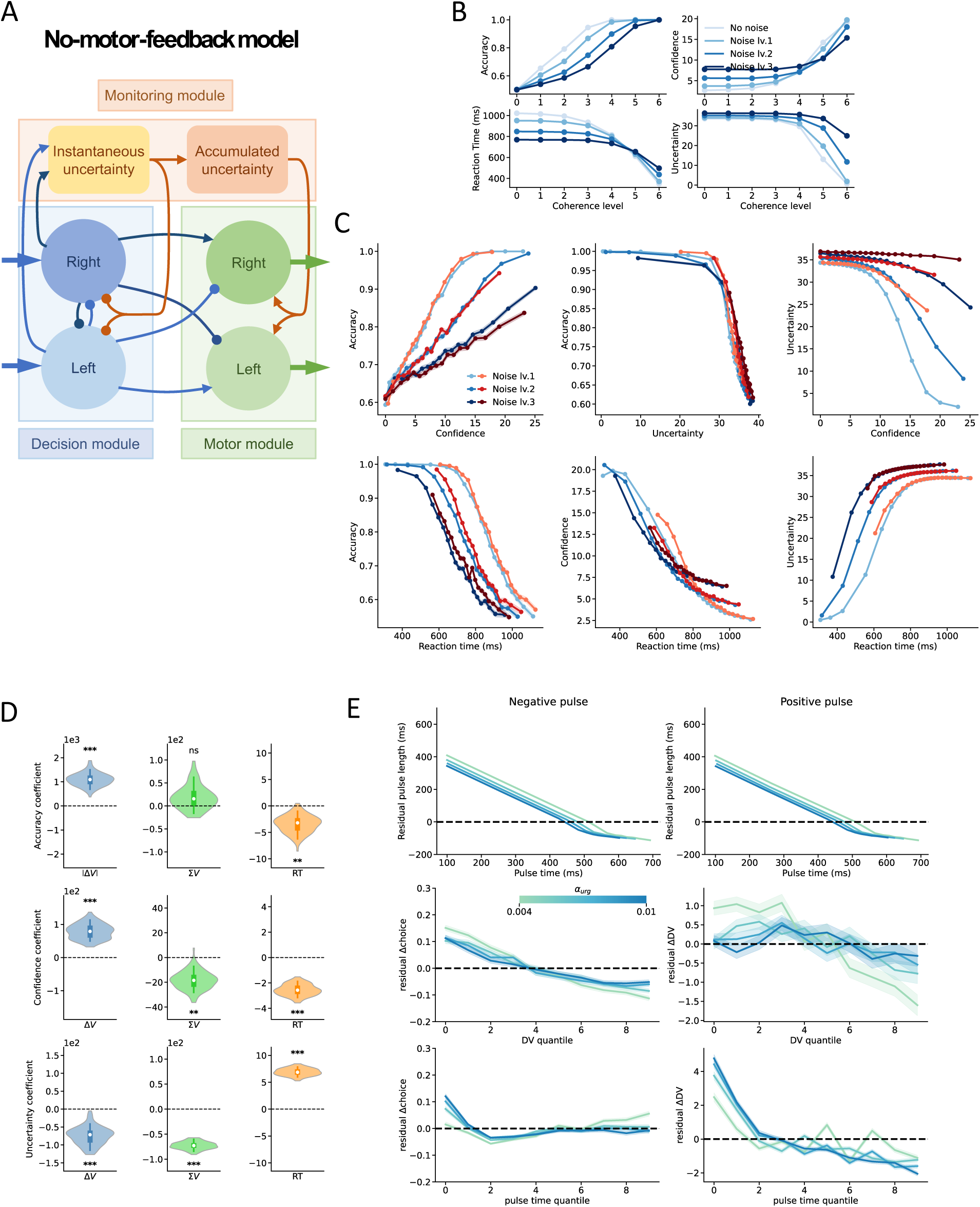
The ‘no-motor-feedback’ reduced model. (**A**) The schematic model that lacks the feedback from the motor module to the decision module, in comparison with the full model. (**B**) The behavioral performance of this reduced model in faces of different noise levels. (**C**) The relationships between accuracy and reaction time with decision uncertainty or decision confidence. (**D**) The dependence of accuracy, confidence and uncertainty on the value difference (*ΔV*), but not the value sum (*ΣV*), reaction time. (**E**) The impacts of stimulus pulses on the choices and decision variables (DVs).

**Figure S6.**
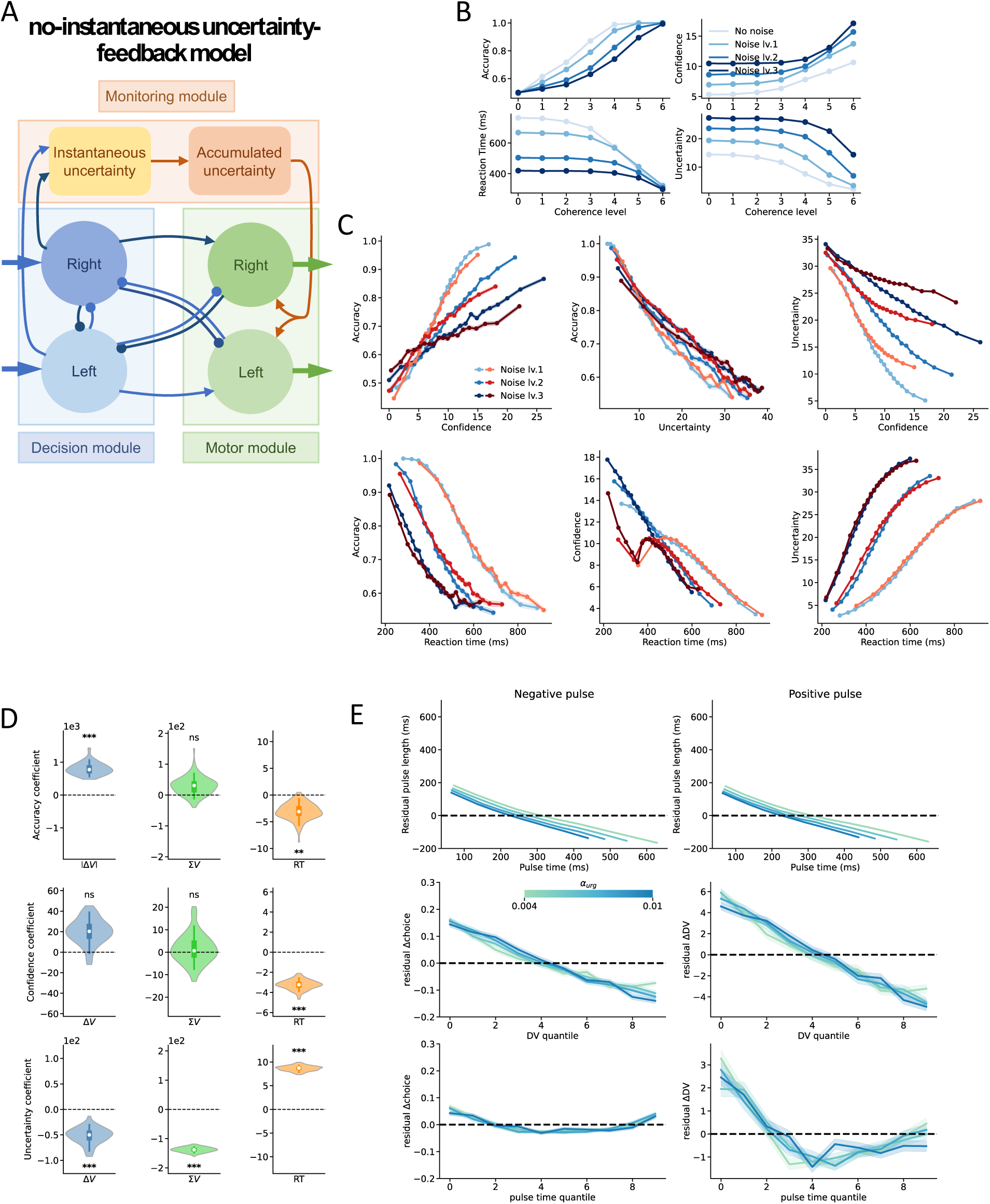
The ‘no-instantaneous uncertainty-feedback’ reduced model. (**A**) The schematic model that lacks the feedback from instantaneous uncertainty module to the decision module, and instead have strong mutual inhibition between decision units. (**B**) The behavioral performance of this reduced model in faces of different noise levels. (**C**) The relationships between accuracy and reaction time with decision uncertainty or decision confidence. (**D**) The dependence of accuracy, confidence and uncertainty on the value difference (*ΔV*), but not the value sum (*ΣV*), reaction time. (**E**) The impacts of stimulus pulses on the choices and decision variables (DVs).

**Figure S7.**
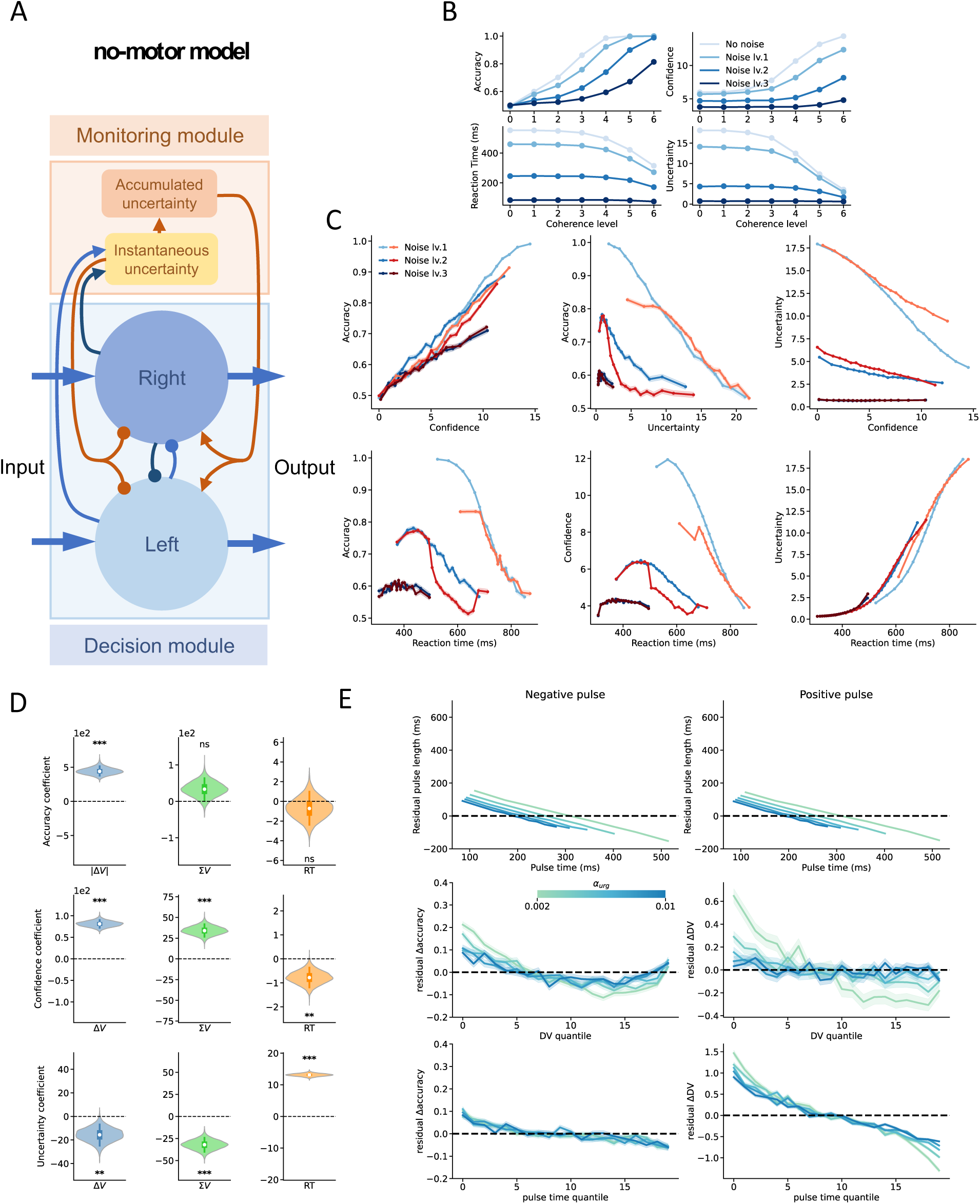
The ‘no-motor’ reduced model. (**A**) The schematic model that lacks the motor module and have strong mutual inhibition between decision units. (**B**) The behavioral performance of this reduced model in faces of different noise levels. (**C**) The relationships between accuracy and reaction time with decision uncertainty or decision confidence. (**D**) The dependence of accuracy, confidence and uncertainty on the value difference (*ΔV*), but not the value sum (*ΣV*), reaction time. (**E**) The impacts of stimulus pulses on the choices and decision variables (DVs).

**Figure S8.**
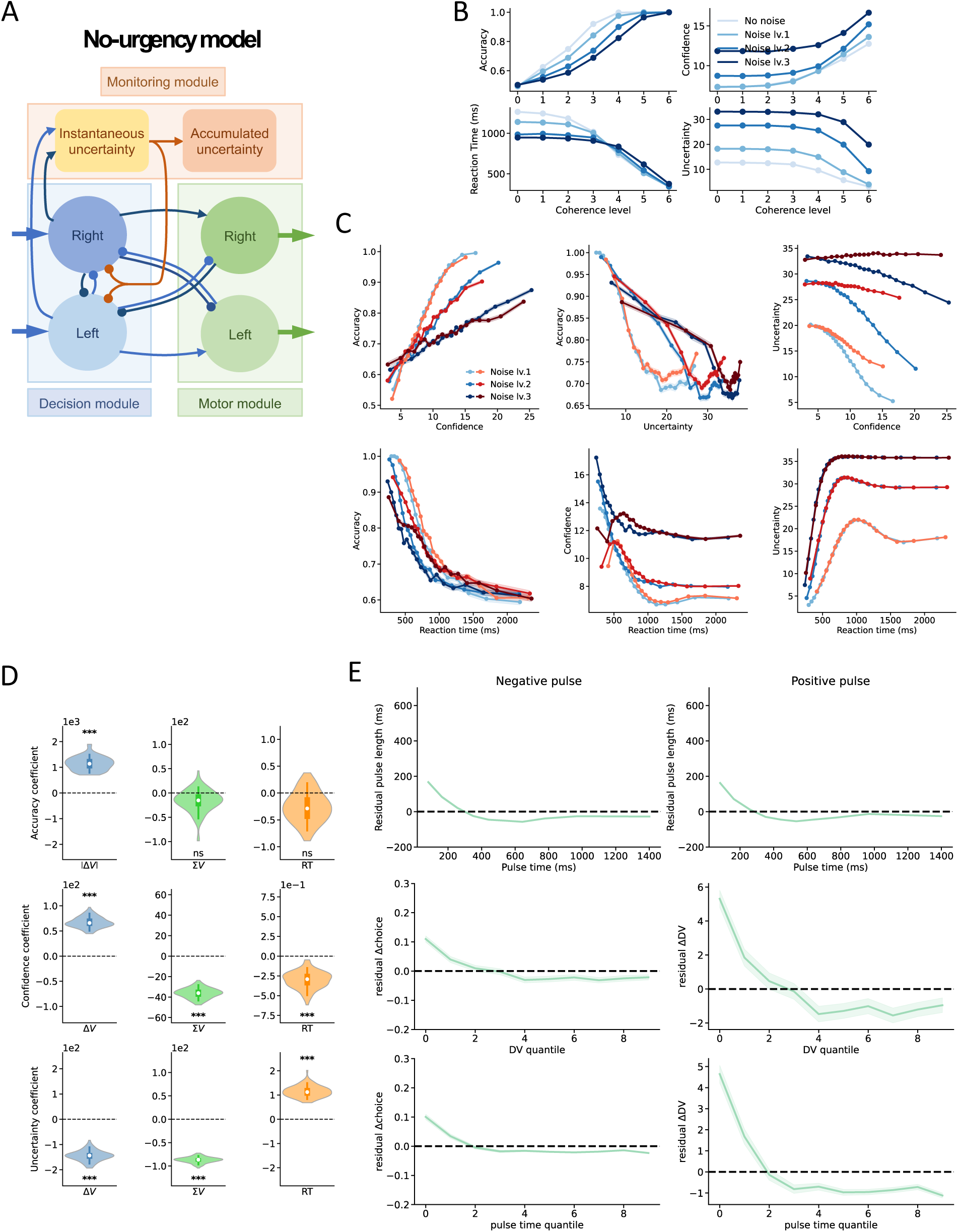
The ‘no-urgency’ reduced model. (**A**) The schematic model that lacks the urgency signal that is fed back from the accumulated uncertainty, and instead has strong mutual inhibition between decision units. (**B**) The behavioral performance of this reduced model in faces of different noise levels. (**C**) The relationships between accuracy and reaction time with decision uncertainty or decision confidence. (**D**) The dependence of accuracy, confidence and uncertainty on the value difference (*ΔV*), but not the value sum (*ΣV*), reaction time. (**E**) The impacts of stimulus pulses on the choices and decision variables (DVs).

**Figure S9.**
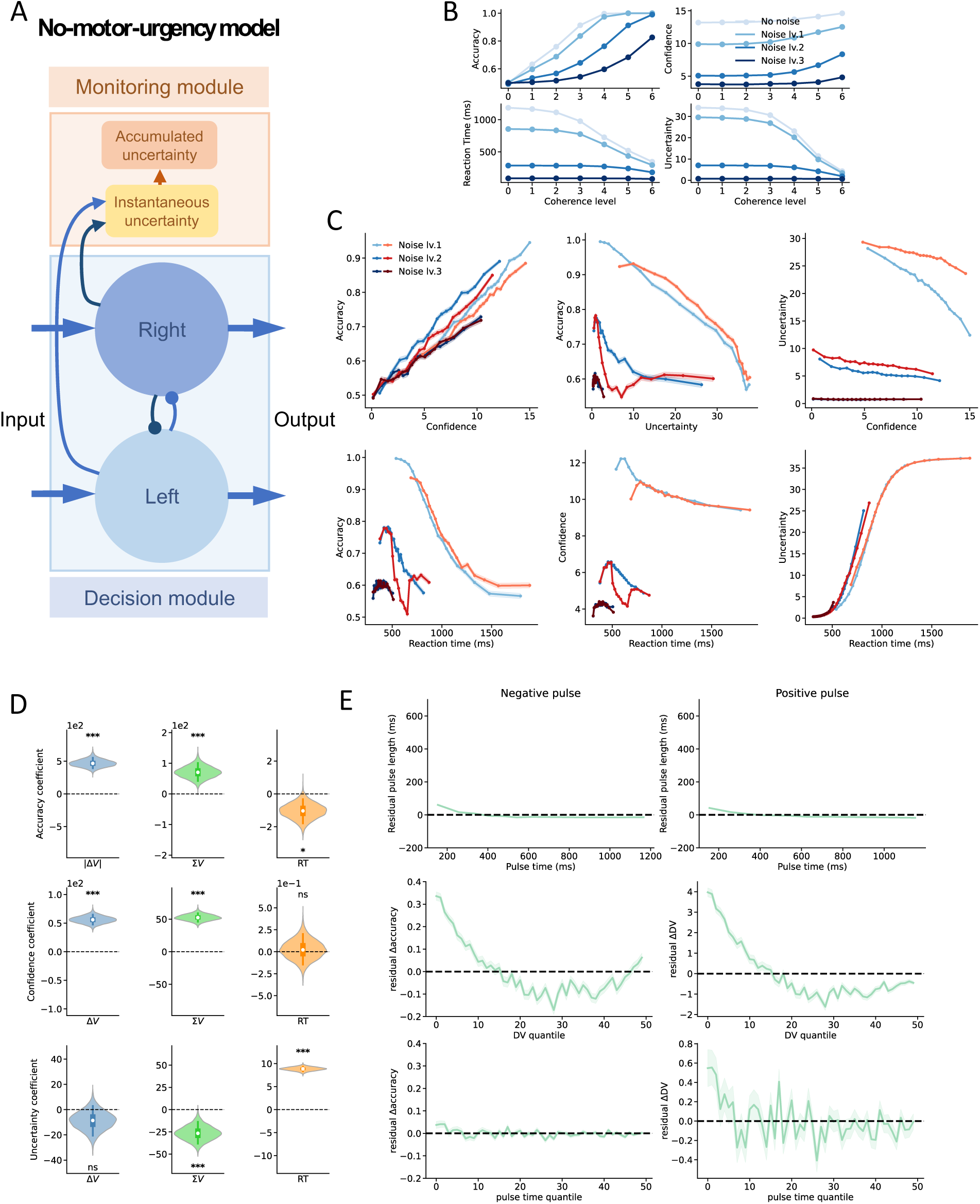
The ‘no-motor-urgency’ reduced model. (**A**) The schematic model that lacks the motor module and feedback from the uncertainty modules, and instead has strong mutual inhibition between decision units. (**B**) The behavioral performance of this reduced model in faces of different noise levels. (**C**) The relationships between accuracy and reaction time with decision uncertainty or decision confidence. (**D**) The dependence of accuracy, confidence and uncertainty on the value difference (*ΔV*), but not the value sum (*ΣV*), reaction time. (**E**) The impacts of stimulus pulses on the choices and decision variables (DVs).

## References

1. L. P. Sugrue, G. S. Corrado, W. T. Newsome, Choosing the greater of two goods: neural currencies for valuation and decision making. Nature Rev. Neurosci. 6, 363–375 (2005).

2. J. I. Gold, M. N. Shadlen, The neural basis of decision making. Annu. Rev. Neurosci. 30, 535–574 (2007).

3. X.-J. Wang, Decision making in recurrent neuronal circuits. Neuron 60, 215–234 (2008).

4. R. Ratcliff, G. McKoon, The diffusion decision model: theory and data for two-choice decision tasks. Neural Comput 20, 873–922 (2008).

5. M. Shinn, N. H. Lam, J. D. Murray, A flexible framework for simulating and fitting generalized drift-diffusion models. eLife 9, e56938 (2020).

6. K. Wong, X.-J. Wang, A Recurrent Network Mechanism of Time Integration in Perceptual Decisions. J. Neurosci. 26, 1314 (2006). doi:10.1523/JNEUROSCI.3733-05.2006.

7. G. Prat-Ortega, K. Wimmer, A. Roxin, J. de la Rocha, Flexible categorization in perceptual decision making. Nature Communications 12, 1283 (2021). doi:10.1038/s41467-021-21501-z.

8. A. Pouget, J. Drugowitsch, A. Kepecs, Confidence and certainty: distinct probabilistic quantities for different goals. Nature Neuroscience 19, 366–374 (2016). doi:10.1038/nn.4240.

9. R. Kiani, M.N. Shadlen, Representation of confidence associated with a decision by neurons in the parietal cortex. Science 324, 759–764 (2009).

10. A. Zylberberg, C. R. Fetsch, M. N. Shadlen, The influence of evidence volatility on choice, reaction time and confidence in a perceptual decision. eLife 5, e17688 (2016). doi:10.7554/eLife.17688.

11. Z. Wei, X.-J. Wang, Confidence estimation as a stochastic process in a eurodynamical system of decision making. J. Neurophysiol. 114, 99–113 (2015).

12. D. Peixoto, J. R. Verhein, R. Kiani, J. C. Kao, P. Nuyujukian, C. Chandrasekaran, J. Brown, S. Fong, S. I. Ryu, K. V. Shenoy, W. T. Newsome, Decoding and perturbing decision states in real time. Nature 591, 604–609 (2021). doi:10.1038/s41586-020-03181-9.

13. T.G. Aquino, J. Cockburn, A.N. Manelak, U. Rutishauser, J.P. O’Doherty, Neurons in human pre-supplementary motor area eoncide key computations for value-based choice. Nature Human Behaviour 7, 970–985 (2023).

14. A. K. Churchland, R. Kiani, M. N. Shadlen, Decision-making with multiple alternatives. Nat. Neurosci. 11, 693–702 (2008).

15. P. Cisek, G. A. Puskas, S. El-Murr, Decisions in changing conditions: the urgency-gating model. J. Neurosci. 29, 11560–11571 (2009).

16. S. Tajima, J. Drugowitsch, A. Pouget, Optimal policy for value-based decision-making. Nat. Commun. 7, 12400 (2016).

17. R. K. Niyogi, K. Wong-Lin, Dynamic excitatory and inhibitory gain modulation can produce flexible, robust and optimal decision-making. PLoS Comput Biol 9, e1003099 (2013). doi:10.1371/journal.pcbi.1003099.

18. P. R. Murphy, E. Boonstra, S. Nieuwenhuis, Global gain modulation generates time-dependent urgency during perceptual choice in humans. Nature Comms. 7, 13526 (2016). DOI: 10.1038/ncomms13526.

19. C. Kunimoto, J. Miller, H. Pashler, Confidence and accuracy of near-threshold discrimination responses. Consciousness and cognition 10, 294–340 (2001).

20. S.M. Fleming, R.S. Weil, Z. Nagy, R.J. Dolan, G. Rees, Relating introspective accuracy to individual differences in brain structure. Science 329, 1541–1543 (2010).

21. E. Rounis, B. Maniscalco, J. C. Rothwell, R. E. Passingham, H. Lau, Theta-burst transcranial magnetic stimulation to the prefrontal cortex impairs metacognitive visual awareness. Cog. Neurosci. 1, 165–175 (2010).

22. J. Su, W. Jia, X. Wan, Task-Specific Neural Representations of Generalizable Metacognitive Control Signals in the Human Dorsal Anterior Cingulate Cortex. J. Neurosci. 42, 1275 (2022). doi:10.1523/JNEUROSCI.1283-21.2021.

23. A. G. Vaccaro, S. M. Fleming, Thinking about thinking: A coordinate-based meta-analysis of neuroimaging studies of metacognitive judgements. Brain and neuroscience advances 2, 2398212818810591 (2018).

24. N. A. A. Atiya, I. Rañó, G. Prasad, K. Wong-Lin, A neural circuit model of decision uncertainty and change-of-mind. Nature Communications 10, 2287 (2019). doi:10.1038/s41467-019-10316-8.

25. N. A. A. Atiya, Q. J. M. Huys, R. J. Dolan, S. M. Fleming, Explaining distortions in metacognition with an attractor network model of decision uncertainty. Plos Comput. Biol. 17, e1009201 (2021).

26. L. Qiu, J. Su, Y. Ni, X. Zhang, Y. Bai, X. Li, X. Wan, The neural system of metacognition accompanying decision-making in the prefrontal cortex. PLOS Biology 16(4) (2018).

27. T. Balsdon, V. Wyart, P. Mamassian, Confidence controls perceptual evidence accumulation. Nature Comms. 11, 1753 (2020). 10.1038/s41467-020-15561-w.

28. X. Li, R. Su, Y. Chen, T. Yang, Optimal policy for uncertainty estimation concurrent with decision making. Cell Reports 42 (2023). doi:10.1016/j.celrep.2023.112232.

29. B. De Martino, S. M. Fleming, N. Garrett, R. J. Dolan, Confidence in value-based choice. Nature Neuroscience 16, 105–110 (2013). 10.1038/nn.3279

30. A. Kepecs, N. Uchida, H.A. Zariwala, Z.F. Mainen, Neural correlates, computation and behavioural impact of decision confidence. Nature 455, 227–231 (2008).

31. R. Kiani, L. Corthell, M. N. Shadlen, Choice certainty is informed by both evidence and decision Time. Neuron 84, 1329–1342 (2014). doi:10.1016/j.neuron.2014.12.015.

32. A.C. Huk, M.N. Shadlen, Neural activity in macaque parietal cortex reflects temporal integration of visual motion signals during perceptual decision making. J. Neurosci. 25, 10420–10436 (2005).

33. A. Zylberberg, P. Barttfeld, M. Sigman, The construction of confidence in a perceptual decision. Front. Integr. Neurosci. 6,79 (2012). 10.3389/fnint.2012.00079

34. T. Hanks, R. Kiani, M. N. Shadlen, A neural mechanism of speed-accuracy tradeoff in macaque area LIP. eLife 3, e02260 (2014). doi:10.7554/eLife.02260.

35. K. Wong, A. C. Huk, M. N. Shalen, X.-J. Wang, Neural circuit dynamics underlying accumulation of time-varying evidence during perceptual decision making. Fron Comput. Neurosci. 2, 1–6 (2007). doi: 10.3389/neuro.10.006.2007.

36. I. Krajbich, C. Armel, A. Rangel, Visual fixations and the computation and comparison of value in simple choice. Nature Neuroscience 13, 1292–1298 (2010). doi:10.1038/nn.2635.

37. G. Tavares, P. Perona, A. Rangel, The Attentional Drift Diffusion Model of Simple Perceptual Decision-Making. Frontiers in Neuroscience 11 (2017). doi:10.3389/fnins.2017.00468.

38. A. Resulaj, R. Kiani, D. M. Wolpert, M. N. Shadlen, Changes of mind in decision-making. Nature 461, 263–266 (2009). doi:10.1038/nature08275.

39. Z. Gao, C. Davis, A. M. Thomas, M. N. Economo, A. M. Abrego, K. Svoboda, C. I. De Zeeuw, N. Li, A cortico-cerebellar loop for motor planning. Nature 563, 113–116 (2018).

40. G. M. Stine, E. M. Trautmann, D. Jeurissen, M. N. Shadlen, A neural mechanism for terminating decisions. Neuron 111, 2601–2613.e5 (2023). doi:10.1016/j.neuron.2023.05.028.

41. A. Koizumi, B. Maniscalco, H. Lau, Does perceptual confidence facilitate cognitive control? Atten. Percept. Psychophys. 77, 1295–1306 (2015). 10.3758/s13414-015-0843-3

42. K. Miyoshi, H. Lau, A decision-congruent heuristic gives superior metacognitive sensitivity under realistic variance assumptions. Psychol. Rev. 127, 655–671 (2020).

43. F. Sun, Y. Ni, W. Lu, S. Wang, J. Su, X. Wan, Confidence bias prescribes neurocomputational mechanism of decision making. Cell Reports (in revision).

44. T. O. Nelson, L. Narens, Metamemory: a theoretical framework and new findings. In the psychology of learning and motivation. Vol. 26. Edited by G. H. Bower, 125–173. New York: Academic Press (1990).

45. M. M. Botvinick, T. S. Braver, C. S. Carter, D. M. Barch, J. D. Cohen. Conflict monitoring and cognitive control. Psychological Review, 108, 624–652 (2001).

46. M. F. S. Rushworth, T. E. J. Behrens. Choice, uncertainty and value in prefrontal and cingulate cortex. Nature Neuroscience, 11, 389–397 (2008).

47. A. Shenhav, M. M. Botvinick, J. D. Cohen. The expected value of control: An integrative theory of anterior cingulate cortex function. Neuron 79, 217–240 (2013).

48. P. Cisek. Cortical mechanisms of action selection: the affordance competition hypothesis. Phil. Trans. Soc. B 362, 1585–1599 (2007).

49. M. A. Carland, D. Thura, P. Cisek. The urge to decide and act: implications for brain function and dysfunction. The Neuroscientist. 25, 491–511 (2019).

50. D. Thura, P. Cisek, Modulation of premotor and primary motor cortical activity during volitional adjustments of speed-accuracy trade-offs. J Neurosci 36, 938–56 (2016).

51. A. Tversky, D. Kahneman, The framing of decisions and the psychology of choice. Science 211, 453–458 (1981).

52. Y. Zhou, D. J. Freedman, Posterior parietal cortex plays a causal role in perceptual and categorical decision. Science 365, 180–185 (2019).

53. L. Zhong, Y. Zhang, C. A. Duan, J. Deng, J. Pan, N.-L. Xu, Causal contributions of parietal cortex to perceptual decision-making during stimulus categorization. Nature Neurosci. 22, 963–973 (2019).

54. J. F. Mejías, X.-J. Wang, Mechanisms of distributed working memory in a large-scale network of macaque neocortex. eLife, 11 (2022).

